# Myeloid lncRNA *LOUP* Mediates Opposing Regulatory Effects of RUNX1 and RUNX1-ETO in t(8;21) AML

**DOI:** 10.1101/2020.07.24.220020

**Authors:** Bon Q. Trinh, Simone Ummarino, Alexander K. Ebralidze, Emiel van der Kouwe, Mahmoud A. Bassal, Tuan M. Nguyen, Rory Coffey, Danielle E. Tenen, Emiliano Fabiani, Carmelo Gurnari, Chan-Shuo Wu, Vladimir Espinosa Angarica, Yanzhou Zhang, Li Ying, Henry Yang, Gerwin Heller, Sisi Chen, Hong Zhang, Abby R. Thurm, Francisco Marchi, Elena Levantini, Philipp B. Staber, Pu Zhang, Maria Teresa Voso, Pier Paolo Pandolfi, Annalisa Di Ruscio, Daniel G. Tenen

## Abstract

The mechanism underlying cell type-specific gene induction conferred by ubiquitous transcription factors as well as disruptions caused by their chimeric derivatives in leukemia is not well understood. Here we investigate whether RNAs coordinate with transcription factors to drive myeloid gene transcription. In an integrated genome-wide approach surveying for gene loci exhibiting concurrent RNA- and DNA-interactions with the broadly expressed transcription factor RUNX1, we identified the long noncoding RNA *LOUP*. This myeloid-specific and polyadenylated lncRNA induces myeloid differentiation and inhibits cell growth, acting as a transcriptional inducer of the myeloid master regulator *PU*.*1*. Mechanistically, *LOUP* recruits RUNX1 to both the *PU*.*1* enhancer and the promoter, leading to the formation of an active chromatin loop. In t(8;21) acute myeloid leukemia, wherein RUNX1 is fused to ETO, the resulting oncogenic fusion protein RUNX1-ETO limits chromatin accessibility at the *LOUP* locus, causing inhibition of *LOUP* and *PU*.*1* expression. These findings highlight the important role of the interplay between cell type-specific RNAs and transcription factors as well as their oncogenic derivatives in modulating lineage-gene activation and raise the possibility that RNA regulators of transcription factors represent alternative targets for therapeutic development.

**KEY POINTS:**

- lncRNA *LOUP* coordinates with RUNX1 to induces *PU*.*1* long-range transcription, conferring myeloid differentiation and inhibiting cell growth.
- RUNX1-ETO limits chromatin accessibility at the *LOUP* locus, causing inhibition of *LOUP* and *PU*.*1* expression in t(8;21) AML.

## INTRODUCTION

Lineage-control genes that dictate cellular identities are often expressed in dynamic and hierarchical patterns.1-3 Disturbance of these established normal patterns results in anomalies.4 In the blood system, the ETS-family transcription factor PU.1 (also known as Spi-1) is essential for myeloid differentiation. *PU*.*1* is silent in most tissues and cell types but expressed at highest levels in myeloid cells including granulocytes and monocytes.^5^ Downregulation of *PU*.*1* impairs myeloid cell differentiation leading to acute myeloid leukemia (AML).^6,7^ *PU*.*1* is a major downstream transcriptional target of Runt-related transcription factor 1 (RUNX1) that is expressed in many different cell types and plays diverse biological roles in hematopoiesis, development of neurons, hair follicles, and skin.^8-12^ In AML with t(8;21) chromosomal translocation, a portion of *RUNX1* containing the Runt DNA binding domain is fused to *ETO*, giving rise to the oncogenic transcription factor fusion RUNX1-ETO.^13,14^ Previously, we have reported that RUNX1-ETO inhibits *PU*.*1* expression^15^ but the mechanism underlying this transcriptional inhibition remains to be determined. In general, how broadly expressed transcription factors, such as RUNX1, modulate cell type- and gene-specific induction and how their chimeric derivatives disrupt this normal regulation in leukemia are poorly understood.

Transcription of many cell type-specific genes are induced by enhancer elements, which are located at variable distances from gene targets.^16,17^ For instance, *PU*.*1* transcription is induced by the formation of a specific chromatin loop resulting from the interaction between the upstream regulatory element (URE) (−17 kb in human and -14 kb in mouse) and the proximal promoter region (PrPr).^18-20^ Interestingly, abrogation of RUNX1-binding motifs at the URE reduces URE-PrPr interaction, resulting in decreased *PU*.*1* expression in myeloid cells.^8,15^ Because RUNX1 is broadly expressed, it remains unclear how this transcription factor modulates chromatin structure in such a gene- and cell type-specific manner.

With advances in whole transcriptome sequencing over the last decade, thousands of noncoding RNAs (ncRNA) have been unveiled.^21^ Arbitrarily defined as ncRNAs having at least 200 nucleotides in length, long noncoding RNAs (lncRNA) are implicated to display tissue-specific expression patterns^22,23^ and might undergo post-transcriptional processing such as splicing and polyadenylation.^24^ Through interactions with DNAs, proteins and other RNAs, lncRNAs regulate fundamental cellular processes including transcription, RNA stability, and DNA methylation.^24-26^ To date, only a few lncRNAs have been precisely mapped and functionally defined,^23^ leaving most lncRNAs poorly annotated and largely unexplored.

In this study, we identified a myeloid-specific lncRNA termed “<u>L</u>ong noncoding RNA <u>O</u>riginating from the <u>U</u>RE of *<u>P</u>U*.*1*”, or *LOUP*, from an integrated genome-wide approach aimed at screening for gene loci exhibiting concurrent RNA- and DNA-interactions with RUNX1. We demonstrated that *LOUP* induces *PU*.*1* expression, conferring myeloid differentiation, and inhibiting cell growth. *LOUP* serves as a central hub in opposing regulation by RUNX1 and its derived oncogenic fusion, RUNX1-ETO. Our findings provide a model explaining how a lineage gene is activated in normal myeloid development and dysregulated in leukemia.

## METHODS

### Cell lines and Cell Culture

U937, HL-60, K562, HEK293T, RAW 264.7, NB4, Jurkat, Kasumi-1 and THP-1 cells were obtained from the American Type Culture Collection (ATCC). U937, HL-60, NB4, Jurkat, and K562 cells were cultured in full RPMI-1640 medium (supplemented with 10% (vol/vol) fetal bovine serum (FBS; Cellgro) and 1% penicillin-streptomycin). Kasumi-1 cells were cultured in the same medium plus 20% (vol/vol) FBS. THP-1 cells were cultured in full RPMI-1640 medium supplemented with 2-mercaptoethanol to a final concentration of 0.05 mM. HEK293T and RAW 264.7 cells were cultured in DMEM supplemented with 10% (vol/vol) FBS and 1% penicillin-streptomycin. All cells were grown at 37°C in 5% (vol/vol) CO2 and humidified incubators.

### AML patient sample collection

Bone Marrow (BM) samples were obtained from newly diagnosed AML patients at the Tor Vergata University Hospital, Rome with informed consent. Diagnoses were performed according to “The 2016 revision to the World Health Organization classification of myeloid neoplasms and acute leukemia”.^27^ Bone marrow mononuclear cells (BM-MNCs) were isolated by Ficoll gradient centrifugation using Lympholyte-H (Cedarlane), according to the manufacturer’s instructions.

Methods for assaying interactions of RNA, DNA, and protein with chromatin, chromatin structure and gene expression manipulation as well as bioinformatic analyses are in supplemental methods.

## RESULTS

### Identification of RUNX1-interacting RNAs at myeloid gene loci

We started out by performing a transcriptome-wide survey for RUNX1-interacting RNAs in the monocytic cell line THP-1 using formaldehyde RNA immunoprecipitation sequencing (RIP-seq).^28,29^ RUNX1-interacting RNAs were captured by anti-RUNX1 antibody (Figures S1A-C) and sequenced by paired-end massively parallel sequencing. By annotating 14,067 high-confident RUNX1-RIP peaks to the GRCh38.p12 gene catalog,^30^ we identified 5,774 gene loci carrying at least one of these peaks (Figure S1D, left). Most of the peaks were detected within transcript bodies and promoters (Figure S1E). To identify genes exhibiting concurrent RUNX1-RNA and RUNX1-DNA interactions, we annotated 24,132 high-confident RUNX1-ChIP peaks to the same gene catalog and identified 13,272 corresponding gene loci (Figure S1D, right). The majority of these peaks were found at intronic, promoter, and intergenic regions (Figure S1F). Because most of peaks identified by both RUNX1-RIP and RUNX1-ChIP peaks were distributed at coding gene loci (Figures 1A-B), we focused our analyses on this gene group. By intersecting these genes with a list of 78 myeloid genes defined by their known roles in myeloid development, or myeloid molecular markers (Table S1), we obtained 15 myeloid gene loci displaying both RUNX1-RIP and RUNX1-ChIP peaks (Figure 1C). *PU*.*1*, a master regulator of myeloid development and a well-known transcriptional target of RUNX1,^8^ was among these genes.

**Figure 1.**
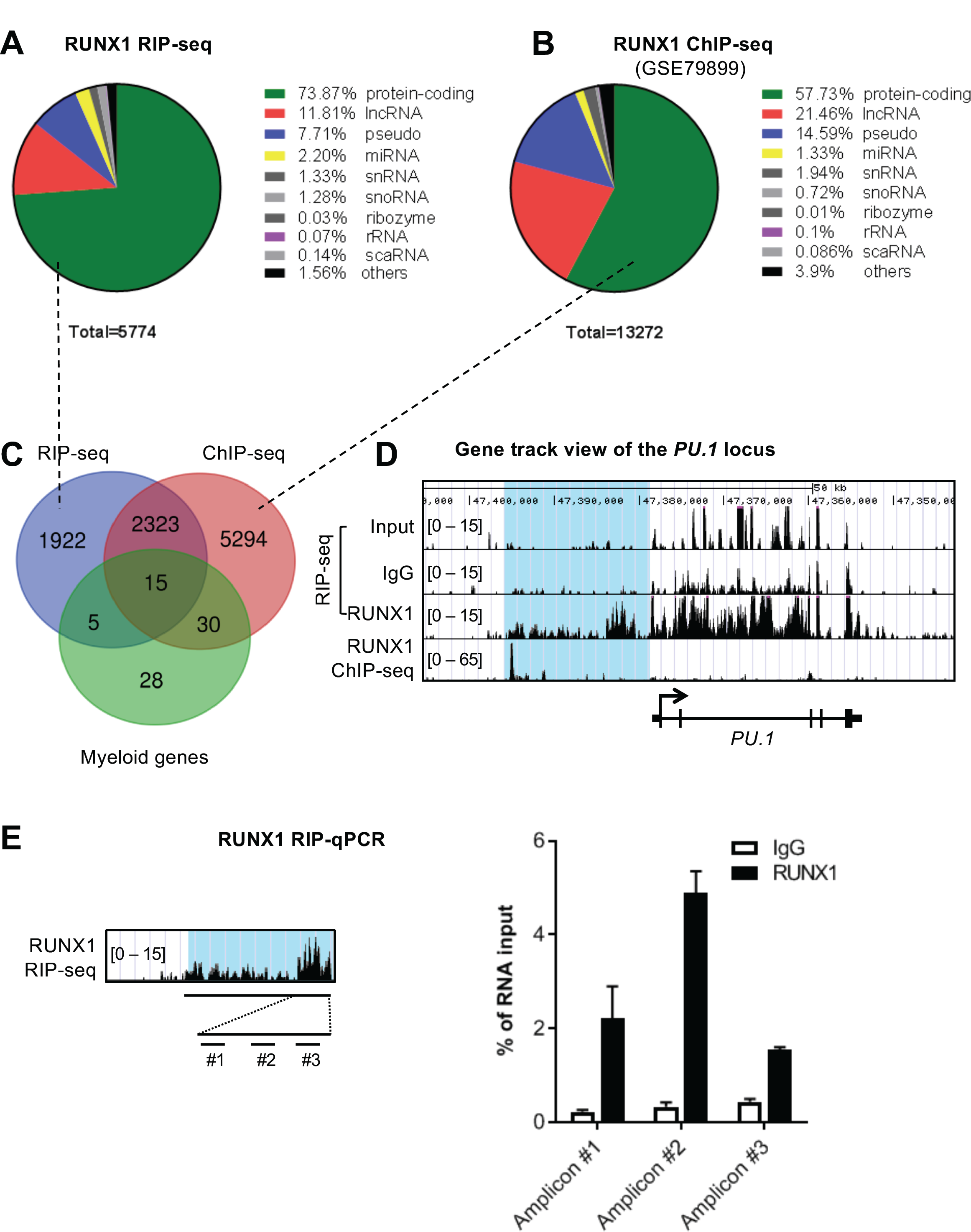
Screening of gene loci exhibiting concurrent RUNX1 RNA and DNA interactions in THP-1 cells. (A and B) Pie charts showing proportions of RUNX1 RIP-seq peaks and RUNX1 ChIP-seq peaks in coding and noncoding gene families. ChIP-seq data was from published source ^59^ under the Gene Expression Omnibus (GEO) accession number: GSE79899. (C) Venn diagram intersecting RUNX1 RIP-seq, RUNX1 ChIP-seq gene lists and the myeloid gene list. (D) Gene track view of the *PU*.*1* locus including the upstream region (highlighted in blue). Shown are RIP-seq tracks (Input, IgG and RUNX1) and RUNX1 ChIP-seq tracks (GSM2108052). Data was integrated in the UCSC genome browser. (E) RUNX1 RIP-qPCR confirmation. Left panel: Location of three PCR amplicons (#1, #2, #3). Right panel: Enrichment of RNAs captured by anti-RUNX1 antibody and IgG control at three amplicons relative to input. Error bars indicate SD (n=3). See also Figure S1 and Table S1.

Intriguingly, we observed RNA peaks at the upstream region of *PU*.*1* (Figure 1D). We further validated this observation by RUNX1 RIP-qPCR (Figure 1E). Additional myeloid genes showing RUNX1-RIP peaks and RUNX1-ChIP peaks are presented in Figure S1G. The presence of previously uncharacterized RNAs, arising from the upstream region of the *PU*.*1* locus, and able to interact with RUNX1, suggests their potential role in controlling *PU*.*1* expression through RUNX1-mediated transcriptional regulation.

### Characterization of the RUNX1-interacting lncRNA *LOUP*

To map the RUNX1-interacting transcript(s), we inspected the RNA expression and epigenetic landscape at the upstream region of the *PU*.*1* locus (Figure 2A). Remarkably, the RNA-seq track view revealed two distinct RNA peaks. A narrow peak was observed at the URE, which corresponded to an area of open chromatin in myeloid cells as indicated by strong DNase I hypersensitivity signals (Figure 2A, DNase-seq). This element was also enriched with histone post-translational modifications such as H3K27ac, H3K4me1, and H3K4me3 (Figure 2A, ChIP-seq), which are typical features of active enhancers.^31,32^ A broad peak was proximal to the promoter region. Notably, these peaks were present in myeloid cell lines (THP-1 and HL-60) and primary monocytes, but not in the lymphoid cell line Jurkat, which does not express *PU*.*1* mRNA, indicating a cell-type specific expression pattern. RT-PCR and Sanger sequencing analysis identified exon junctions connecting these two peaks in both human and murine cell lines (Figure S2A). Strand-specific RT-PCR analysis confirmed that the transcript is sense with respect to the *PU*.*1* gene (Figure 2B). To locate the 5’ end, we inspected Cap analysis gene expression sequencing (CAGE-seq) tracks from the FANTOM5 project,^33^ and identified a strong CAGE-seq peak, located within the URE and in the sense genomic orientation (Figure 2A, CAGE-seq), suggesting the presence of the 5’ end of a transcript. Using the P5-linker ligation method outlined in Figure S2B, we identified the 5’ end including a transcription start site (TSS) of the RNA within the homology region 1 (H1) of the URE^18^ (Figure S2C). Although a splicing event was detected within the second exon, intron retention was dominant as shown by the presence of a ∼2.3 Kb major transcript and a ∼1.0 Kb minor transcript (Figures 2C and S2D). The transcripts were detectable in the myeloid cell line U937, but not in the lymphoid cell line Jurkat, further indicating their cell-type specificity (Figure 2C). Notably, the RNA exhibited very low coding potential similar to that of other known lncRNAs (Figure S2E) as assessed by PhyloCSF software.^34^ Additionally, no known protein domains were found (data not shown) using PFAM software.^35^ Thus, we named the RNA transcript “long noncoding RNA <u>o</u>riginating from the <u>U</u>RE of *<u>P</u>U*.*1*”, or *LOUP*. qRT-PCR analyses of subcellular fractionations revealed that *LOUP* resides in both the cytoplasm and the nucleoplasm compartments, and was particularly enriched in the chromatin fraction (Figure S2F). The lncRNA is polyadenylated, being detected from oligo(dT)-primed cDNAs (Figure 2B) and enriched in the polyA^+^ RNA fraction (Figures 2C-D and S2G). *LOUP* is low abundant lncRNA; the spliced form is expressed as ∼40, 14, and 5 copies per cells in U937, HL-60, and NB4, respectively (Figure 2E). The lncRNA was barely detectable as its premature (non-spliced) form in total RNA as well as in the nuclear RNA fraction (Figures S2H-I). Altogether, these findings established *LOUP* as a polyadenylated lncRNA that emanates from the URE and extends toward the PrPr.

**Figure 2.**
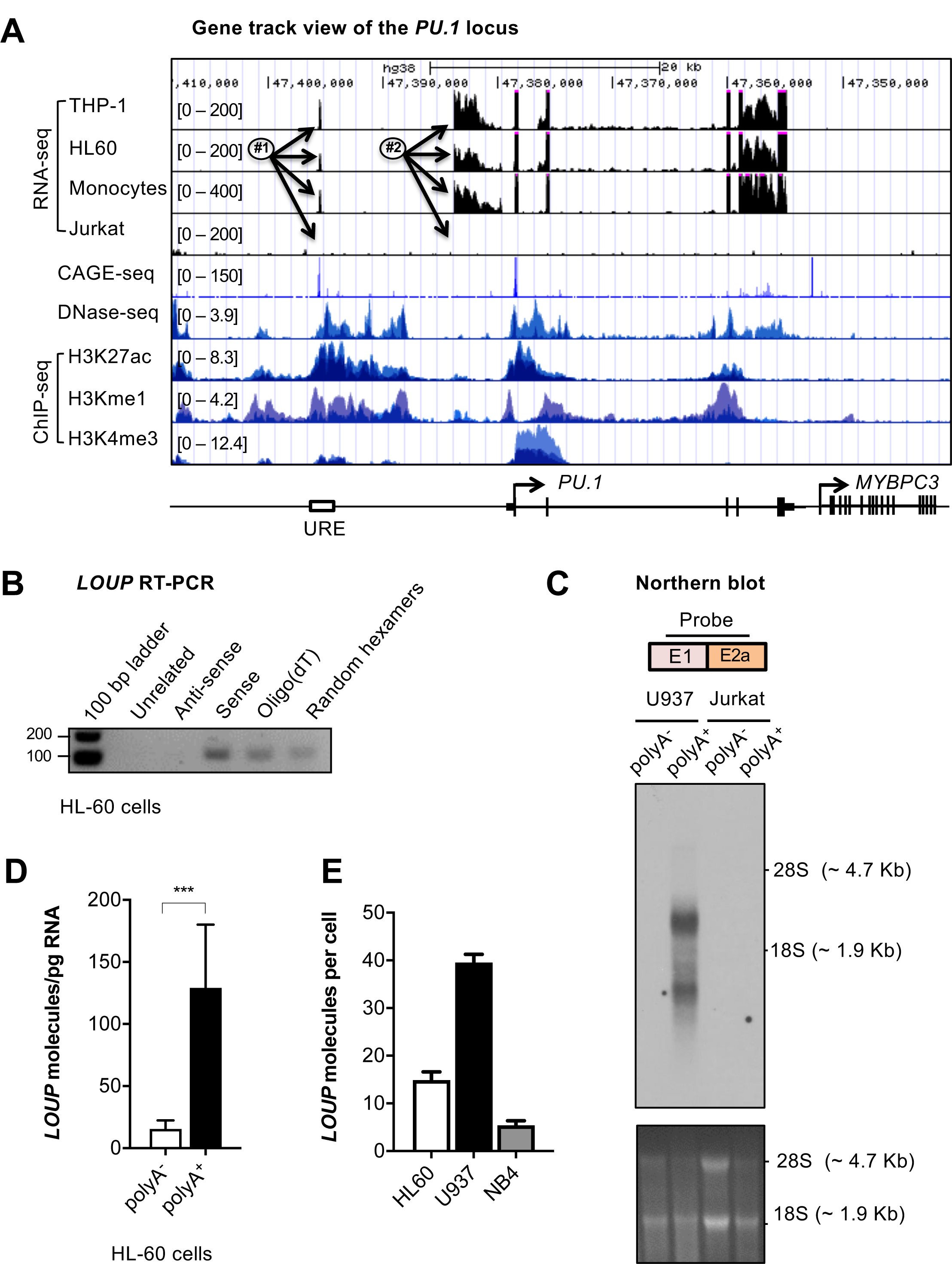
Characterization of long noncoding RNA *LOUP*. (A) Gene track view of the genomic region encompassing the *PU*.*1* locus. RNA-seq tracks include THP-1, HL60, primary monocytes, and Jurkat. DNAse-seq and ChIP-seq are overlay tracks of monocyte and myeloid cell lines. These data were processed from published data in GEO (see methods for details). CAGE-seq track was imported from the FANTOM5 project. #1, #2 and arrows point to locations of the RNA peaks. (B) RT-PCR analysis of *LOUP*’s transcript features. First-strand cDNAs were generated from HL-60 total RNA using a primer that does not anneal to the *PU*.*1* locus (Unrelated), Random hexamers, Oligo(dT), and strand-specific primers (Anti-sense and Sense). (C) Northern blot analysis of *LOUP*. polyA^-^ and polyA^+^ RNA fractions were isolated from U937 and Jurkat cells. Top panel: schematic of the probe location spanning exon junction (E1 and E2a; see Figure S2D). Middle panel: Northern blot detection of *LOUP*’s major and minor transcripts. Lower panel: RNA gel demonstrating relative migration between 28S and 18S rRNAs stained with ethidium bromide. (D) qRT-PCR analysis of *LOUP* levels in polyA^-^ and polyA^+^ RNA fractions isolated from HL-60 cells. Error bars indicate SD (n=3). ***p < 0.001. (E) Calculation of *LOUP* transcript per cell by qRT-PCR. *LOUP* RNA standard curve was generated by *in vitro* transcription. Error bars indicate SD (n=3). See also Figure S2.

### *LOUP* is myeloid-specific lncRNA that is co-expressed with myeloid lineage gene *PU*.*1*

We sought to explore *LOUP* expression in normal tissues and cell types. By examining the *LOUP* transcript profile in different human tissue types from the Illumina Body Map dataset, we noticed that this lncRNA was barely detectable in most tissues but elevated in leukocytes (Figure 3A). Remarkably, comparing with two of its closest neighbor genes, *PU*.*1* and *SLC39A13* (Figure S2D), the *LOUP* expression pattern was similar to that of *PU*.*1* mRNA (Figures 3A-B) but not of *SLC39A13* (Figure S3A). Additionally, *LOUP* transcript levels were not correlated with that of its interacting partner, *RUNX1* (Figure S3B). To further delineate the relationship between *LOUP* and *PU*.*1* transcript levels and lineage identity in individual blood cells, we employed single-cell RNA-seq analyses (scRNA-seq). scRNA-seq data of human mononuclear cells isolated from peripheral blood (PBMC) and bone marrow (BMMC) were retrieved from the 10x Genomic Project^36^ and pooled together to maximize coverage of hematopoietic cell lineages (Figure S3C). Notably, *LOUP* and *PU*.*1* were both enriched in the myeloid cells, comprising of monocytes, macrophages and granulocytes (Figures S3D-E). Expectedly, *RUNX1* was broadly expressed in myeloid cells as well as lymphoid cells (T, B, and Natural Killer (NK)) (Figure S3F). By stratifying the mononuclear cell population into *LOUP*^high^/*PU*.*1*^high^ and *LOUP*^low^/*PU*.*1*^low^ groups based on *LOUP* and *PU*.*1* expression levels (see methods for details), we noted that *LOUP*^low^/*PU*.*1*^low^ cells were associated with T, B, and NK cells. Remarkably, 99.3% of *LOUP*^high^/*PU*.*1*^high^ cells were linked to the myeloid identity (Figure 3C). Consistent with this observation, top biological processes associated with expression of *LOUP* and *PU*.*1* were mono/macrophage and granulocyte functions (Figure S3G and Table S2). We further examined expression patterns of *LOUP* and *PU*.*1* during myeloid differentiation. qRT-PCR analyses of purified murine hematopoietic cell populations showed low *Loup* levels in long-term hematopoietic stem cells (LT-HSC), short-term hematopoietic stem cells (ST-HSC), common myeloid progenitors (CMP), and megakaryocyte-erythroid progenitors (MEP).

**Figure 3.**
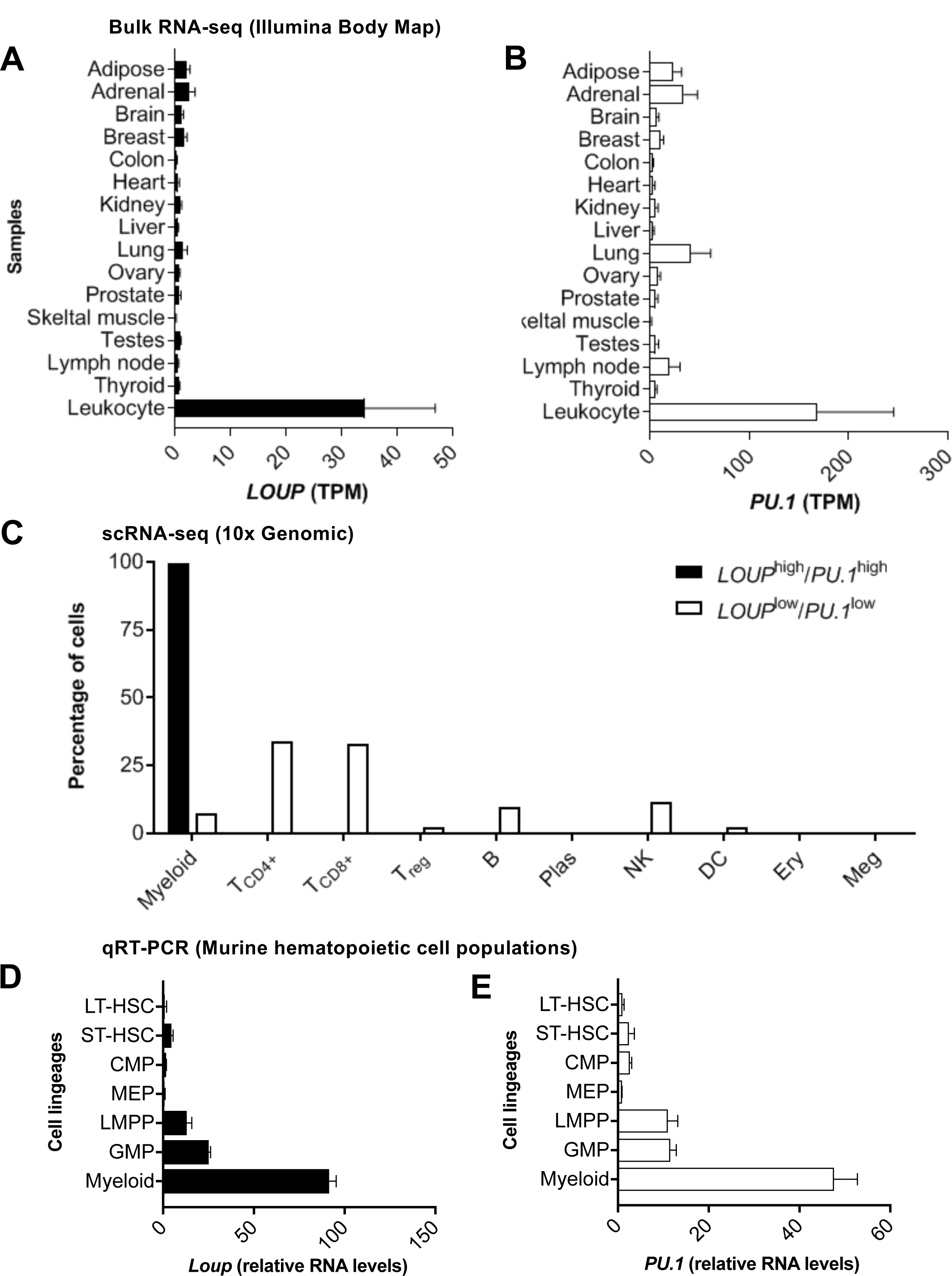
Expression profiles of *LOUP* and *PU*.
*1* in normal tissues and cell lineages. (A and B) Transcript profiles of *LOUP* and *PU*.*1* in human tissues. Shown are transcript counts from the Illumina Body Map RNA-seq data dataset (AEArrayExpress: E-MTAB-513). Error bars indicate SD (n=2). (C) Proportion of cell lineages corresponding to *LOUP* and *PU*.*1* transcript levels. Myeloid: includes monocytes, macrophages and granulocytes; T_CD4+_: T helper cell; T_CD8+:_ Cytotoxic T cell; T_reg_: Regulatory T cell; B: B lymphocyte; Plas: Plasma cell; NK: Natural killer cell; DC: Dendritic cell; Ery: Erythrocyte; Meg: Megakaryocyte. (D and E) qRT-PCR analysis of *Loup* RNA and *PU*.*1* mRNA levels in murine hematopoietic stem, progenitor, and mature (myeloid) cell populations. LT-HSC: long-term hematopoietic stem cells; ST-HSC: short-term hematopoietic stem cells; CMP: common myeloid progenitors, MEP: megakaryocyte-erythroid progenitors; LMPP: lymphoid-primed multipotent progenitors; GMP: granulocyte-macrophage progenitors, myeloid cells (Mac1^+^Gr1^+^). Data are shown relative to LT-HSC. Error bars indicate SD (n=2). See also Figure S3 and Table S2.

Remarkably, *Loup* expression was elevated in myeloid progenitor cells (granulocyte-macrophage progenitors, GMP) and was highest in definitive myeloid cells (Figure 3D). A similar expression pattern was seen with *PU*.*1* (Figures 3E). Taken together, our data indicated that *LOUP* and *PU*.*1* transcript levels were associated with the myeloid identity, warranting further investigation regarding molecular relationship between *LOUP* and *PU*.*1* in myeloid cells.

### *LOUP* induces *PU*.*1* expression, promotes myeloid differentiation and inhibits cell growth

To test our hypothesis that *LOUP* induces *PU*.*1* expression, we investigated the impact of loss-of-function of *LOUP* on *PU*.*1* expression. In order to deplete *LOUP*, we employed CRISPR/Cas9 genome-editing platform which introduces small insertion and deletion (indel) mutations in the *LOUP* gene via the non-homologous end-joining (NHEJ) DNA repair mechanism.^37,38^ The macrophage cell line U937 that expresses high levels of *LOUP* (Figure 2E) was stably transduced with lentiviruses carrying Cas9 and *LOUP*-targeting or non-targeting sgRNAs. Double-positive mCherry (CAS9) and eGFP (sgRNA) cells were selected by fluorescence-activated cell sorting (FACS) (Figures 4A and S4A), and derived cell clones were analyzed by Sanger DNA sequencing and Inference of CRISPR edits (ICE) analysis.^39^ Cell clones having indels at *LOUP*-targeting genomic locations (Figures S4B-D) displayed >80% depletion of *LOUP* RNA levels (Figure 4B, left panel). This depletion was paralleled by a significant reduction in *PU*.*1* mRNA levels (Figure 4B, right panel). In gain-of-function experiments, transient *in trans*-overexpression of *LOUP* in K562 cells resulted in significant induction of *PU*.*1* (Figure 4C).

**Figure 4.**
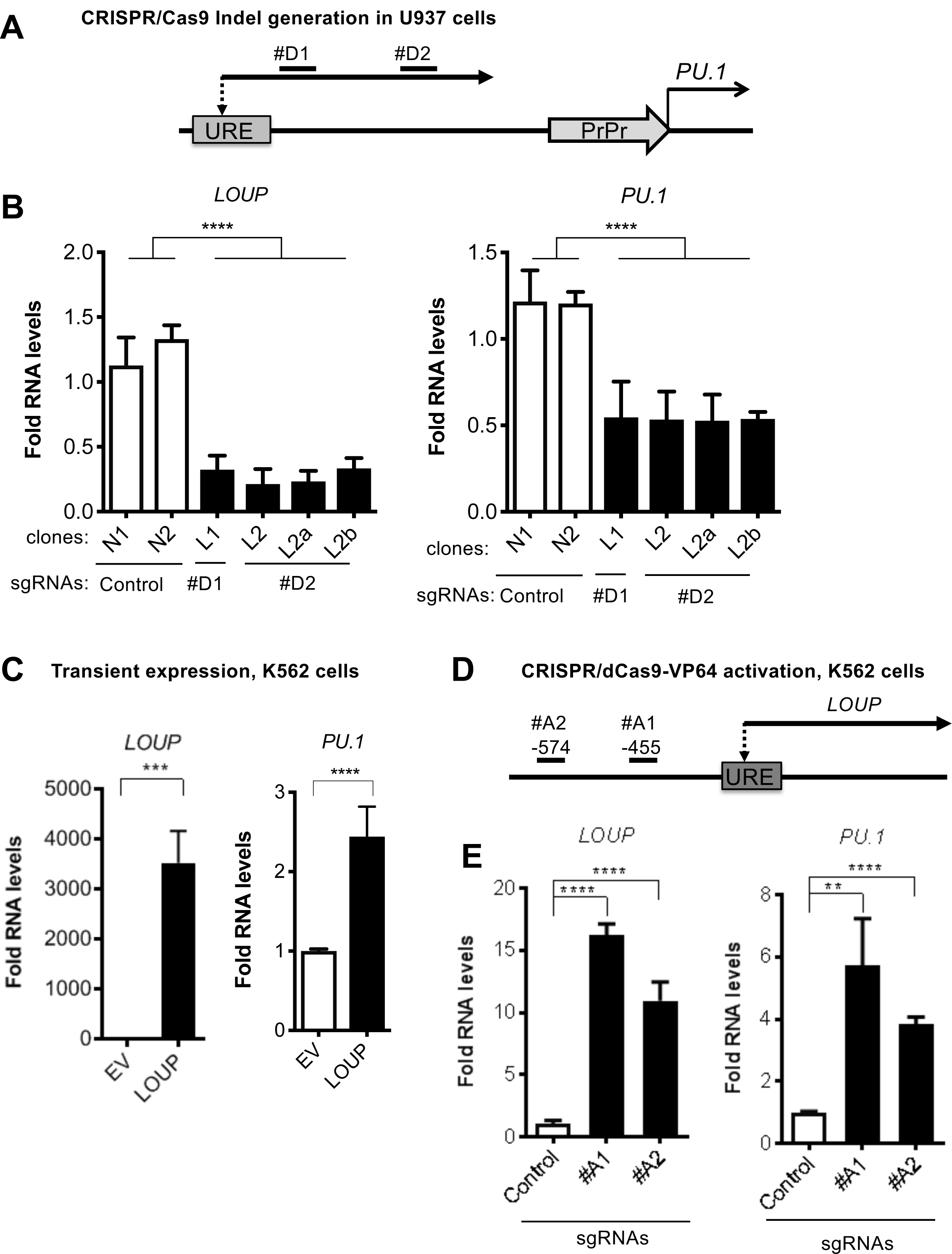
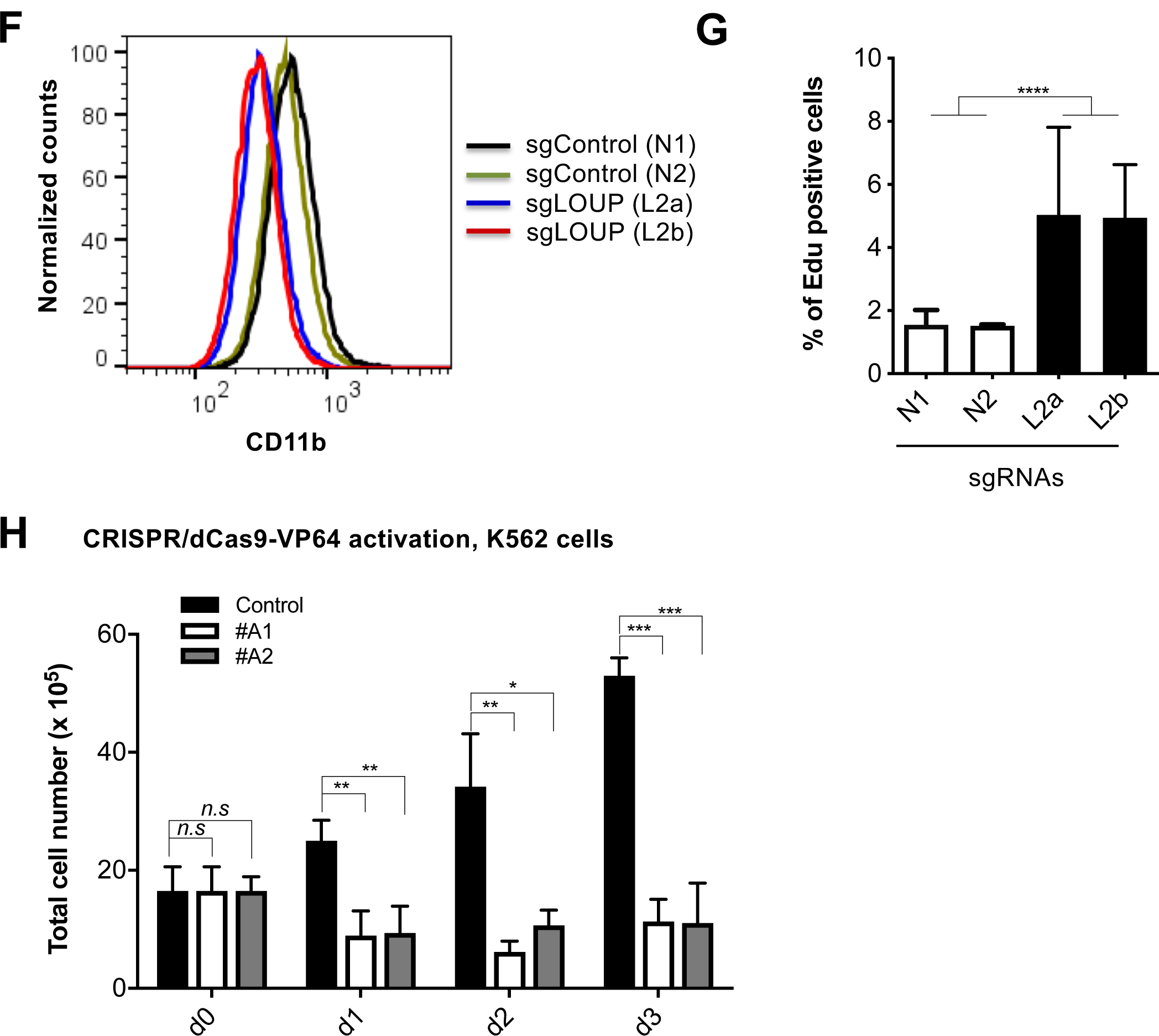
The effect of *LOUP* on *PU*.*1* expression, myeloid differentiation and cell growth. (A) Schematic diagram of the upstream genomic region of the *PU*.*1* locus. Shown are sgRNA-binding sites (#D1 and #D2) for *LOUP* depletion using CRISPR/Cas9 technology. (B) qRT-PCR expression analysis for *LOUP* (left panel) and *PU*.*1* (right panel) in non-targeting (N) and *LOUP*-targeting (L) U937 cell clones. Data are shown relative to N1 control. (C) qRT-PCR expression analysis of *LOUP* RNA (left panel) and *PU*.*1* mRNA (right panel) in K562 cells transfected with *LOUP* cDNA or empty vector (EV) by electroporation. (D) Schematic diagram of the *LOUP* promoter region showing sgRNA-binding sites (#A1 and #A2) for *LOUP* induction. Distance from the TSS of *LOUP* is indicated in bp (D) qRT-PCR expression analysis of *LOUP* (left panel) and of *PU*.*1* (right panel) in K562 dCas9-VP64-stable cells infected with *LOUP*-targeting (#A1 and #A2) or non-targeting (control) sgRNAs. (E) FACS analysis of CD11b myeloid marker in U937 cell clones with *LOUP* homozygous indels (L2a and L2b) and controls (N1 and N2) using PACBLUE-conjugated CD11b antibody. (F) Edu incorporation was measured by flow cytometry for cell proliferation. (G) Trypan blue exclusion and manual cell counts for kinetics of cell growth (shControl *v*.*s*. shLOUP (#A1 and #A2). Error bars indicate SD (n=3). \**p* < 0.05; \*\**p* < 0.01; \*\*\**p* < 0.001, \*\*\*\**p* < 0.0001, *n*.*s*: not significant. See also Figure S4.

Remarkably, *in cis* locus-specific induction of endogenous *LOUP* via the CRISPR/dCas9-VP64 activation system yielded a comparable increase in *PU*.*1* expression, despite producing lower *LOUP* transcripts than the ectopic *in trans*-expression (Figures 4D-E). Consistent with the important role of *PU*.*1* in myeloid differentiation,^6,7,40,41^ *LOUP* depletion was associated with a reduction in expression of the myeloid marker CD11b (Figure 4F). Furthermore, *LOUP* depletion increases cell proliferation whereas enforced *LOUP* reduced cell number suggesting that *LOUP* inhibits cell growth (Figures 4G-H). Together, these results demonstrate that *LOUP* promotes myeloid differentiation and inhibits cell growth and that this lncRNA regulator exerts its inducing effect on *PU*.*1* expression primarily in cis.

### *LOUP* induces enhancer-promoter communication by interacting with chromatin at the *PU*.*1* locus

We have previously reported that the formation of a chromatin loop mediated by URE-PrPr interaction is crucial for *PU*.*1* induction.^18,19^ Because *LOUP* arises from the URE and extends toward the PrPr, we reasoned that *LOUP* drives long-range transcription of *PU*.*1* by promoting URE-PrPr interaction. To elucidate this, we quantified interaction strengths of the URE with the PrPr and the surrounding area by chromosome conformation capture and Taqman qPCR (3C-qPCR) (Figure 5A). Consistent with previous reports,^18,19^ we detected strong interaction between the URE and the PrPr, but not between the URE and other genomic regions, including the upstream *PU*.*1* promoter, intergenic sequences, and the *MYBPC3* gene body downstream of the *PU*.*1* locus. Interestingly, *LOUP* depletion caused a significant reduction in URE-PrPr communication (Figures 5B). To provide evidence supporting our prediction that *LOUP* recruits the URE to the PrPr by physically interacting with the two elements, we employed the Chromatin Isolation by RNA Purification (ChIRP) assay.^42^ Biotinylated *LOUP*-tiling oligos were able to capture endogenous *LOUP* RNA in U937 cells (Figure 5C). Enrichment of the URE and the PrPr co-captured with *LOUP* RNA was observed in ChIRPed samples with *LOUP*-tiling probes but not LacZ-tiling controls, suggesting that *LOUP* occupies both the URE and the PrPr (Figure 5D). Taken together, our data indicate that by interacting and bringing to close proximity two regulatory elements, the URE and the PrPr, *LOUP* promotes the formation of a functional chromatin loop within the *PU*.*1* locus that is critical in inducing *PU*.*1* expression.

**Figure 5.**
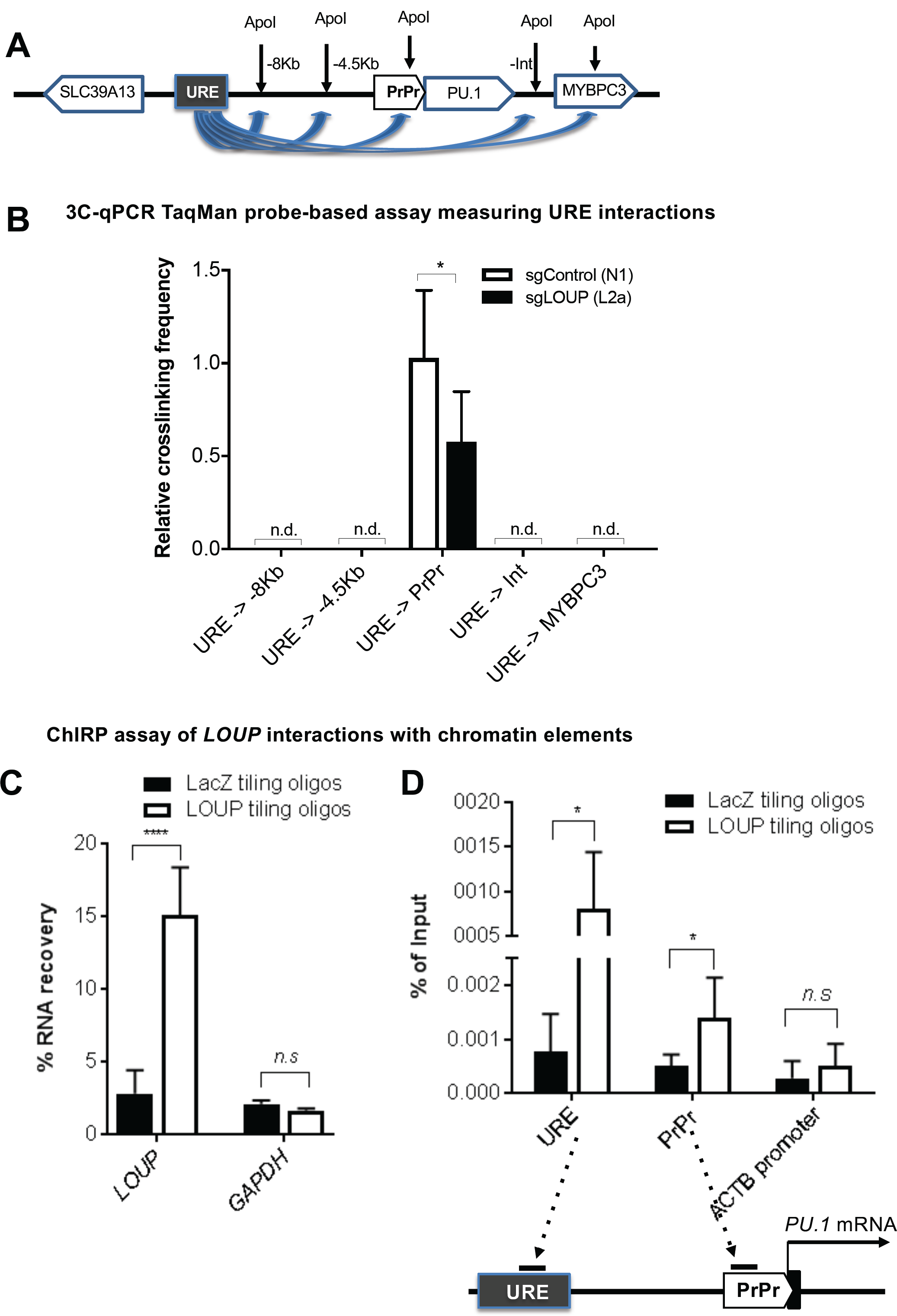
3C and ChIRP assays measuring the effect of *LOUP* on chromatin looping. (A) Schematic diagram illustrating potential 3C interactions between the URE and genomic viewpoints surrounding the *PU*.*1* locus. Included are restriction recognition sites of ApoI used in the assay. -8Kb, and -4Kb: distances from the PrPr in kilo bases. Int: intergenic. (B) 3C-qPCR TaqMan probe-based assay comparing crosslinking frequencies at chromatin viewpoints. The U937 cell clone L2a, carrying a *LOUP* homozygous indel that does not alter the recognition pattern of ApoI (Figure S4D), was used to compare with non-targeting control (sgControl, N1). n.d.: not detectable. (C) qRT-PCR assay evaluating levels of *LOUP* RNA and control *GAPDH* captured by biotinylated *LOUP*-tiling and LacZ-tiling probes using ChIRP. (D) ChIRP assay assessing *LOUP* occupancies at the URE, the PrPr, and *ACTB* promoter. LOUP-tiling oligos were used to capture endogenous *LOUP* in U937 cells. LacZ-tiling oligos were used as negative control. Error bars indicate SD (n=3); \**p* < 0.05; \*\*\*\**p* < 0.0001, *n*.*s*: not significant.

### *LOUP* binds the Runt domain of RUNX1 and coordinates recruitment of RUNX1 to the enhancer and the promoter

We next sought to gain a deeper mechanistic understanding of how *LOUP* modulates the chromatin structure in a gene specific manner. Point mutations abolishing the RUNX binding sites in the URE are known to disrupt chromosomal interactions between the URE and the PrPr.^15^ Additionally, we demonstrated that *LOUP* interacts with RUNX1 at the *PU*.*1* locus (Figure 1). Therefore, we asked whether *LOUP* mediates the URE-PrPr interaction by cooperating with RUNX1. In line with the previous finding in murine cells,^15^ we observed RUNX1 occupancy at the URE in primary CD34^+^ cells isolated from healthy donors and patients with AML. Importantly, we noticed a peak at the PrPr, indicating that RUNX1 also occupies the PrPr (Figure 6A). We further inspected the genomic region surrounding the PrPr and found a RUNX-DNA binding consensus motif at -220 bp relative to the *PU*.*1* mRNA transcription start site. To determine if this motif is functional, we performed biotinylated DNA pull-down (DNAP) assays. Wild-type probes, containing the RUNX consensus motifs embedded in the URE and the PrPr, efficiently captured endogenous RUNX1 from U937 nuclear extracts. In contrast, probes mutating the RUNX1 binding sequence, displayed drastic reductions in RUNX1 occupancy (Figures 6B and S5A). These results suggest that RUNX1 binds its DNA consensus motif at both the URE and the PrPr. RUNX1 is known to form homodimers to modulate transcription.^43,44^ Thus, we reasoned that *LOUP* promotes looping formation by conferring occupancy of RUNX1 dimers concurrently at their binding motifs within the URE and the PrPr. Indeed, *LOUP* depletion reduced RUNX1 occupancy at both the URE and the PrPr (Figure 6C), indicating that *LOUP* promotes placement of RUNX1 dimers at the URE and the PrPr. By aligning *LOUP* sequence with itself using the Basic Local Alignment Search Tool (BLAST), we unexpectedly uncovered a highly repetitive region (RR) of 670 bp near the 3’ end of *LOUP* (Figure S5B). Interestingly, by performing RNA pull-down assays (RNAP) assay, we noted that biotinylated *LOUP* RR was able to capture endogenous RUNX1 proteins in U937 nuclear extracts at a level that was comparable to biotinylated full-length *LOUP*, indicating that the RR contains RUNX1-binding region (Figure 6D). To further locate the binding region, we first computed potential interaction strength of putative elements within the RR to RUNX1 protein by using the catRAPID algorithm.^45^ By doing so, we identified two ∼100 bp candidate regions, termed region 1 (R1) and region 2 (R2) within, with high interaction scores (Figures S5C and 6E). RNAP analysis confirmed that R1 and R2 bind recombinant RUNX1 (Figure 6F). Additionally, recombinant Runt domain of RUNX1 was able to bind R1 and R2 (Figure 6G), suggesting that the Runt domain is responsible for *LOUP* binding. These data, together, suggests that *LOUP* binds RUNX1 and coordinates deposition of RUNX1 dimers to the URE and the PrPr.

**Figure 6.**
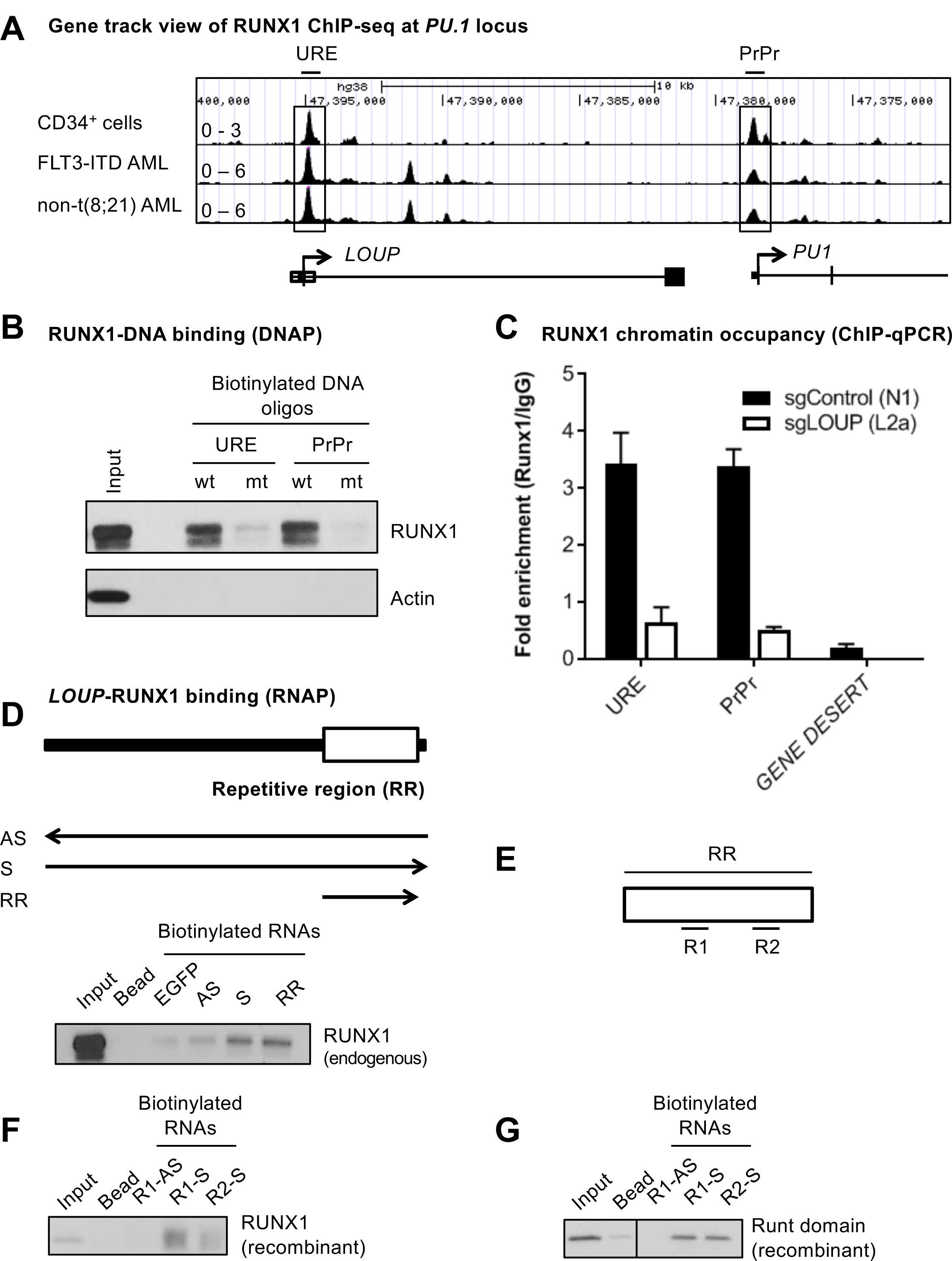
*LOUP* cooperates with RUNX1 to facilitate URE-PrPr interaction. (A) Gene track view of the ∼26 kb region encompassing the URE and the PrPr. Shown are RUNX1 ChIP-seq tracks derived from CD34^+^ cells from healthy donors (GSM1097884), an AML patient with FLT3-ITD AML (GSM1581788), and a non-t(8;21) AML patient (GSM722708) (top panel). The bottom panel is a schematic showing the corresponding genomic locations of *LOUP* and the 5’ region of *PU*.*1*. (B) DNA pull-down assay showing binding of RUNX1 to the RUNX1-binding motifs at the URE and the PrPr. Proteins captured by biotinylated DNA oligos (wt: wildtype oligo containing RUNX1-binding motif, mt: oligo with mutated RUNX1-binding motif) in U937 nuclear lysate were detected by immunoblot. (C) ChIP-qPCR analysis of RUNX1 occupancy at the URE and the PrPr. *LOUP*-depleted U937 (sgLOUP, L2a) and control (sgControl, N1) clones were used. PCR amplicons include the URE (contains known RUNX1-binding motif at the URE), PrPr (contains putative RUNX1-binding motif in the PrPr) and GENE DESERT (a genome region that is devoid of protein-coding genes). Error bars indicate SD (n=3). (D) RNA pull-down analysis of the RUNX1-*LOUP* interaction. Upper panel: Schematic diagram of *LOUP* showing relative position of the repetitive region RR. Arrows underneath the diagram illustrate direction and relative lengths of *in vitro*-transcribed and biotin-labeled *LOUP* fragments (Bead: no RNA control; EGFP: *EGFP* mRNA control; AS: full-length antisense control; S: full-length sense, and RR: repetitive region). Lower panel: *LOUP* fragments were incubated with U937 nuclear lysate. Retrieved proteins were identified by immunoblot. (E) Schematic diagram of the repetitive region RR showing predicted binding regions R1 and R2. (F and G) RNAP binding analysis of R1 and R2 with recombinant full-length and Runt domain of RUNX1. *In vitro*-transcribed and biotin-labeled RNAs include R1-AS (R1 antisense control); R1-S (R1 sense); and R2-S (R2 sense). The vertical line demarcates where an unrelated lane was removed from the figure. See also Figure S5.

### RUNX1-ETO down-regulates *LOUP* in t(8;21) AML by inhibiting histone H3 acetylation and reducing promoter accessibility

We further examined how the oncogenic fusion protein RUNX1-ETO, derived from t(8;21) chromosomal translocation, affects the regulatory function of *LOUP*. By examining *LOUP* transcript profiles in an AML RNA-seq dataset downloaded from The Cancer Genome Atlas (TCGA), we noticed that *LOUP* RNA levels were significantly lower in t(8;21) AML patients as compared to AML patients with normal karyotype (Figure 7A, left panel). Consistent with our data demonstrating the *PU*.*1* is a downstream target of *LOUP, PU*.*1* levels were also lower in t(8;21) AML patients (Figure 7A, right panel). These finding were further confirmed by qRT-PCR using patient samples (Figure 7B). Thus, we reasoned that *LOUP* may act as an inhibitory target of RUNX1-ETO in t(8;21) AML. Indeed, depletion of RUNX1-ETO in t(8;21) AML cells Kasumi-1, resulted in a robust increase in *LOUP* transcript levels which was accompanied by a significant induction in *PU*.*1* mRNA (Figure 7C). RUNX1-ETO is capable of recruiting Nuclear Receptor Corepressor Histone Deacetylase Complex and associates with histone deacetylase activity.^46-48^ To examine whether RUNX1-ETO inhibits *LOUP* transcription by affecting local histone acetylation, we analyzed histone acetylation and chromatin accessibility at the URE, where *LOUP* transcription is initiated, upon depletion of RUNX1-ETO.^49^ As expected, knockdown of RUNX1-ETO reduces RUNX1-ETO occupancy at the URE (Figure 7D, top panel). Interestingly, depletion of RUNX1-ETO resulted in robust induction of the H3K9Ac histone acetylation mark that is associated with active promoters and Dnase I accessibility at the URE (Figure 7D, middle and bottom panels), indicating that RUNX1-ETO inhibits *LOUP* transcription by deacetylating and limiting its promoter accessibility.

**Figure 7.**
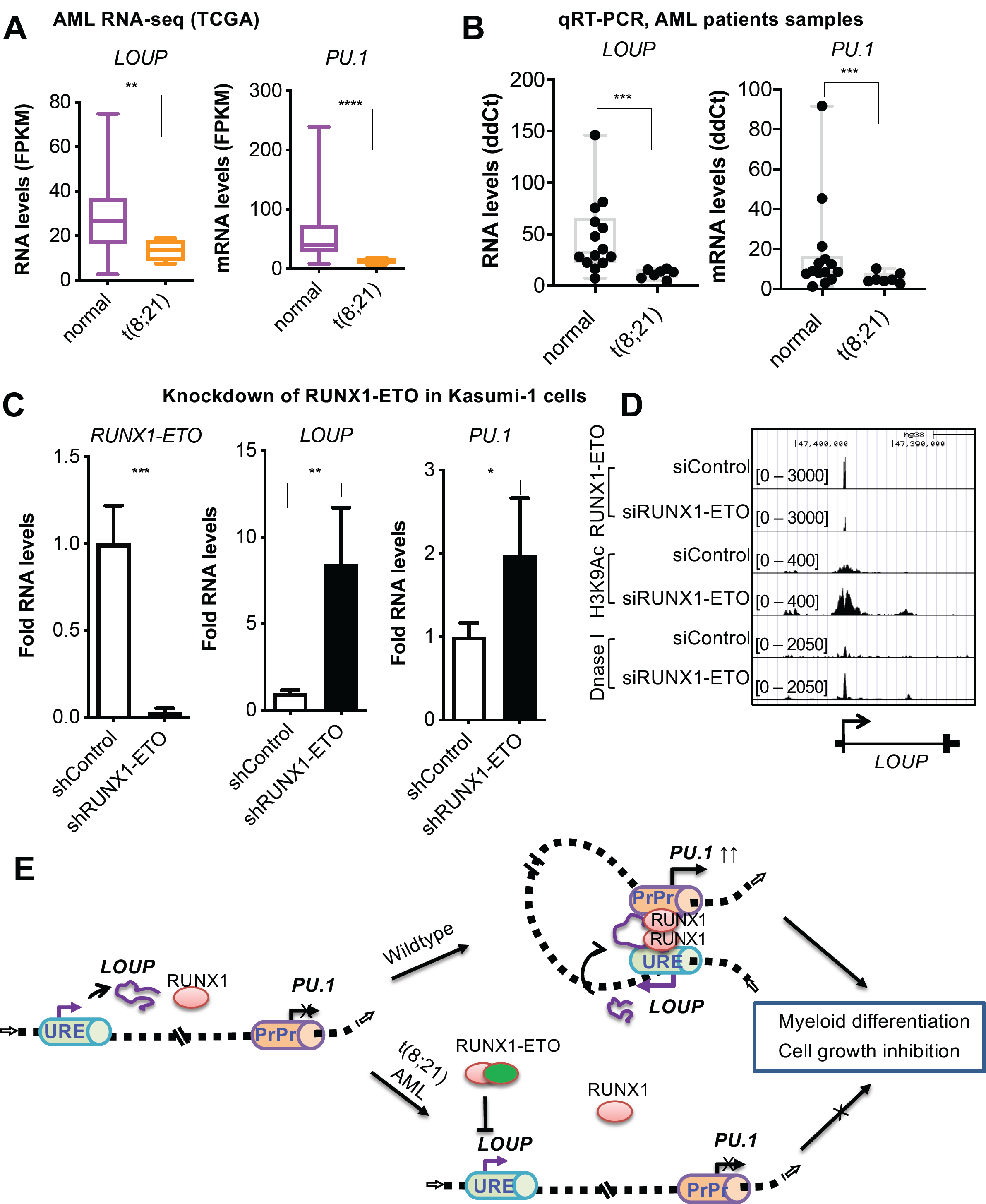
Effects of RUNX1-ETO on regulatory function of *LOUP*. (A) Transcript count for *LOUP* levels in AML patient samples (RNA-seq data was retrieved from TCGA portal. normal: normal karyotype n=87, t(8;21) n=7); Mann–Whitney U test: **p<0.01, ****p<0.0001. (B) RT-qPCR analysis of AML patient samples. normal: normal karyotype (n=14), t(8;21) (n=7). Mann–Whitney U test: ***p<0.001. (C) qRT-PCR expression analysis of *RUNX1-ETO* (left panel), *LOUP* RNA (middle panel) and *PU*.*1* mRNA (right panel) in Kasumi-1 cells transfected with Renilla-targeting shRNA (shControl) and RUNX1-ETO targeting shRNA (shRUNX1-ETO). Error bars indicate SD (n=4), *p<0.05, **p<0.01, ***p<0.001. (D) Gene track view at the *LOUP* locus including the URE where *LOUP* transcription initiation is located. Shown are RUNX1-ETO ChIP-seq tracks (top panels), H3K9Ac ChIP-seq tracks (middle panels) and DNase-seq tracks of Kasumi-1 cells upon depletion of RUNX1-ETO. Cells were transfected with either nontargeting siRNA (siControl) or RUNX1-ETO-targerting siRNA. Data was processed from published dataset (GEO: GSE29222) and integrated in the UCSC genome browser. (E) Model of how *LOUP* coordinates with RUNX1 to modulate chromatin looping resulting in PU.1 induction, myeloid differentiation, and cell growth, and how RUNX1-ETO interferes with *LOUP*-mediated molecular functions.

In summary, we established *LOUP* as a myeloid-specific lncRNA that promotes myeloid differentiation and inhibits cell growth via cooperating with RUNX1 to induce *PU*.*1* expression, and that RUNX1-ETO disrupts the action of *LOUP* in t(8;21) AML. Thus, lncRNA *LOUP* acts as a regulatory hub delivering opposing effects from a broadly expressed transcription factor and its oncogenic derivative on long-range transcription of an important lineage gene (Figure 7E).

## DISCUSSION

In this study, we discovered that RUNX1, which is expressed and exerts its regulatory roles in diverse cell types,^50,51^ cooperates with a myeloid-specific lncRNA *LOUP* to induce long-range transcription of *PU*.*1*, and that RUNX1-ETO impairs *LOUP*-mediated *PU*.*1* induction by inhibiting *LOUP* expression in t(8;21) AML. Our study reported several important mechanistic findings. We reveal *LOUP* as a cellular RNA-interacting partner of RUNX1. We also demonstrate that *LOUP* recruits RUNX1 to respective RUNX1-binding motifs at both the URE and the PrPr, thereby promoting formation of the URE-PrPr chromatin loop at the *PU*.*1* locus. Additionally, we identify a repetitive region serving as the RUNX1-binding platform for *LOUP*. Furthermore, we show that *LOUP* is a inhibitory target of RUNX1-ETO, in t(8;21) AML. These findings provide important insight into how long-range transcription is induced in a gene-specific manner by ubiquitous transcription factors and how their chimeric derivative disrupt normal gene induction in leukemia.

Our findings that RUNX1, known to be crucial for the URE-PrPr interaction, occupies both the URE and the PrPr of the *PU*.*1* locus, provides a molecular understanding of locus-specific activation. We propose that, once the URE and the PrPr are brought into close proximity, RUNX1 molecules that are parts of separate URE- and PrPr-bound complexes might interact, resulting in the formation of the URE-PrPr (enhancer-promoter) transcriptional activation complex. In supporting of this mechanism, RUNX1 sites at enhancers and promoters have been shown to be critical for induction of *CSF2* (encoding GM-CSF), *CD34*, and *CEBPA* (encoding C/EBPα),^43,52-54^ suggesting that RUNX1 could also contribute to specific enhancer-promoter docking at these gene loci. In line with this notion, locus-specific enhancer-promoter interaction could be induced by artificially tethering transcription factor to promoter.^55^ Our findings, therefore, support a model in which specific and on-target enhancer-promoter interactions are achieved by transcription factors, bound to specific motifs both at the enhancer and the target promoter, that are able to dimerize or multimerize, thereby helping to fuse enhancer and promoter transcriptional complexes together.

How chromatin-bound protein complexes at enhancers and target promoters are brought together in a highly specific manner is still poorly understood. Our findings offer several exciting avenues that might explain how locus-specific induction is accomplished. First, we demonstrated that *LOUP* modulates recruitment of RUNX1 to its binding motifs at both the URE and the PrPr, suggesting that *LOUP* might serve as an “RNA bridge”, bringing the separate RUNX1-containing-URE and –PrPr transcriptional complexes into proximity which finally fused into an URE-PrPr complex via RUNX1 dimerization. Second, locus specificity might also be enhanced based on our finding that *LOUP* arises from the URE and acts *in cis* to modulate chromatin looping at the nearby *PU*.*1* locus. Accordingly, even when a small number of transcripts are being produced, local molecular concentration of *LOUP* could be enriched enough to profoundly influence rapid *PU*.*1* mRNA induction. Indeed, we found that *LOUP* is a low-abundance lncRNA but is enriched in the chromatin fraction. Third, we revealed that *LOUP* is expressed exclusively in myeloid cells. This could explain why RUNX1, which is expressed in diverse cell types, induces URE-PrPr interaction and *PU*.*1* expression specifically in myeloid cells.

These findings, together, provide mechanistic understanding of gene-specific enhancer-promoter interaction and cell type-specific gene induction.

Our findings also contribute to the growing body of knowledge with regard to molecular functions of lncRNAs. Indeed, among thousands of lncRNAs that are implicated to arise throughout the genome, only a few have been precisely mapped and molecularly characterized.^23^ The herein described lncRNA *LOUP*, presenting as spliced and polyadenylated transcripts, binds the Runt domain of RUNX1 via a repetitive region. To our knowledge, *LOUP* is the first cellular RNA-interacting partner of RUNX1 being reported. Remarkably, we also discovered that *LOUP* is down-regulated by RUNX1-ETO. It also worth mentioning that a normal allele of *RUNX1* is retained alongside RUNX1-ETO fusion gene in t(8;21) AML cells^56^ and that RUNX1-ETO is implicated to exert opposing effect by competing with RUNX1 for binding to protein partners and the same chromatin locations.^49,57,58^ Collectively, our findings uncover a heretofore-unknown cross-regulation and molecular interactions of lncRNAs with transcription factors and their oncogenic derivatives, providing mechanistic understanding underlying their molecular functions.

In summary, we identified lncRNA *LOUP* with several important molecular features, including cell-type specific expression and harboring a RUNX1-binding platform enabling *LOUP* to coordinate with RUNX1 to drive long-range transcription of *PU*.*1* in myeloid cells. *LOUP*, a downstream inhibited target of the oncogenic fusion protein RUNX1-ETO, is capable of inducing myeloid differentiation and inhibiting cell growth. Our finding raises the possibility that RNA regulators of transcription factor represent alternative targets for therapeutic development and provide a molecular mechanism explaining, at least in part, how ubiquitous transcription factors contribute to enhancer-promoter communication in both cell-type and gene-specific manner and how their chimeric derivatives disrupt this normal regulation in leukemia.

## ACKNOWLEDGEMENTS

This work was supported by the following grants and awards. K01CA222707 to BQT; R50 CA211304 to AKE; NCI R00 CA188595 and the Italian Association for Cancer Research (AIRC) awards to ADR; AIRC 5×1000 call “Metastatic disease: the key unmet need in oncology” to MYNERVA project, #21267 to MTV; NCI R35 CA197697, P01HL131477, the Singapore Ministry of Health’s National Medical Research Council under its Singapore Translational Research (STaR) Investigator Award, and by the National Research Foundation Singapore and the Singapore Ministry of Education under its Research Centres of Excellence initiative to DGT. BQT thanks Linus Tsai and Touati Benoukraf for technical advice. The authors thank the Iannis Aifantis Lab for the generous gift of the sgRNA cloning vector, and Susumu Kobayashi, Robert Welner, To-Ha Thai, Constanze Bonnifer, and Li Chai for insightful comments. We also thank Junyan Zhang, Qiling Zhou and all the members of the Tenen Laboratory for technical assistance and helpful suggestions.

## AUTHORSHIP CONTRIBUTIONS

DGT and BQT designed the study with contribution from AKE, PBS, MTV, PPP, and ADR. BQT, SU, AKE, SC, PZ, HZ, EL, FM, EK, EF, CG and ART performed experiments. BQT, VEA and GH analyzed ChIP-seq data; BQT and MB analyzed RIP-seq and scRNA-seq data; BQT and TMN analyzed bulk RNA-seq data and performed CRISPRa; RC and DET designed and performed RIP-seq experiment; LY, HY and BQT analyzed TCGA data, CW and BQT performed PhyloCSF analysis; BQT and SU drew schematics; BQT and DGT wrote the manuscript with input from authors, especially, AKE, EL, MB, TMN, PBS, MTV, PPP, and ADR. DGT supervised the project.

