## Supplemental methods and data for "Myeloid lncRNA *LOUP* Mediates Opposing Regulatory Effects of RUNX1 and RUNX1-ETO in t(8;21) AML"

[1]Harvard Medical School Initiative for RNA Medicine, Harvard Medical School, Boston, MA, 02115, USA ;[2]Division of Endocrinology, Diabetes, and Metabolism, Beth Israel Deaconess Medical Center, Boston, MA, USA; [3]Cancer Science Institute, National University of Singapore, 117599, Singapore [4]Harvard Stem Cell Institute, Harvard Medical School, Boston, MA, 02115, USA; [5]University of California Los Angeles, CA, 90095, USA; [6]University of Florida, FL, 32611, USA; [7]Institute of Biomedical Technologies, National Research Council (CNR), Area della Ricerca di Pisa, Pisa, 56124, Italy; [8]Department of Medicine I, Division of Hematology, Medical University of Vienna, Vienna, Austria [9]University of Rome Tor Vergata, Roma, 00133, Italy; [10]University of Eastern Piedmont, Department of Translational Medicine, Novara, 28100, Italy

[§]Corresponding Author and Lead Contact:
Daniel G. Tenen


Bon Q. Trinh


### SUPPLEMENTAL METHODOLOGY

#### Lentiviral generation

Lentiviral particles were generated following our optimized protocol [1]. In brief, HEK293T cells were plated overnight to reach 80-85% confluency on the next day. Cells were then co-transfected with the viral expression vector plus packaging plasmids (pMD2.G and psPAX2, Addgene) using Lipofectamine 2000 (Life Technologies). At 48 h and 72 h thereafter, culture supernatants were collected and filtered through a 0.45-mm PVDF filter (Millipore). Viruses were further concentrated using PEG-it® Virus Precipitation Solution (System Biosciences).

**Plasmid generation**

*LOUP* cDNA is in pCMV-SPORT6 plasmid (Dharmacon). Short hairpin RNAs (shRNA) targeting Renilla (shControl) and RUNX1-ETO (shRUNX1-ETO) were cloned into lentiviral vector containing GFP in an optimized 'miRE' context [2]. shRNA sequences are provided in Table S3.

**Generation of *LOUP*-depleted U937 cells (CRISPR/Cas9)**

FUCas9Cherry [3] (Addgene) was used as expression vector to generate mCherry-Cas9 lentiviral particles as described above. U937 cells were transduced with these particles using the TransDux® reagent (System Biosciences). Cas9-stable cells were then selected by several rounds of FACS sorting for mCherry positivity. *LOUP*-targeting sgRNAs were designed using Cas-Designer [4] and cloned into the pLVx U6se EF1a sfPac vector which carries eGFP (kind gift from Dr. Iannis Aifantis). To avoid disruption of the URE, known to be critical for *PU.1* induction (Li et al. 2001), we designed single-guide RNAs (sgRNA) targeting two distinct regions of the *LOUP* gene: (1) the *LOUP* intronic area downstream of the URE, and (2) the intronic area immidiately upstream of the second exon of the *LOUP* gene (~ 15 kb downstream from the URE). Cas9-stable cells were then transduced with eGFP-sgRNA lentiviruses. Cells expressing high levels of both eGFP and mCherry were FACS sorted, one cell per well, into 96-well plates. Genomic DNA from cell clones were isolated using DNeasy Blood & Tissue Kit kit (QIAGEN) and used for PCR amplifying CRISPR/Cas9 target sites. PCR products were sequenced and indel profile were analyzed by Inference of CRISPR edits (ICE) software [5]. Cell clones having homozygous indels were verified by Sanger sequencing. Primer and sgRNA sequences are provided in Table S3.

**Generation of CRISPR activation cells (CRISPRa)**

sgRNAs targeting the region 500 bp upstream of the *LOUP* transcriptional start site were designed using Cas-Designer [4]. The sgRNAs were then cloned into the pXR502 plasmid as previously described [6]. K562 cells stably expressing dCas9-VP64 were generated via lentiviral delivery of dCas9-VP64-Blast [7] and Blasticidin selection. dCas9-VP64 stable cells were transduced with lentiviruses that packaged the sgRNA-cloned pXR502 plasmids as previously described [6]. One-day post-transduction, cells were selected with puromycin for 2-3 days before collection for analysis. sgRNA sequences are provided in Table S3.

**Plasmid transfections**

K562 cells, in exponential growth, were electroporated with expression plasmids using program T16, kit V (Lonza). Electroporated cells were incubated at 37°C overnight in a 5% $CO_2$ incubator. The next day, cells were changed to fresh medium. Cells were harvested at 48 h after electroporation.

**Cellular fractionation, RNA extraction, RT-PCR and qPCR analysis**

Cultured cells were washed with phosphate-buffered saline (PBS). Total RNA was extracted with Trizol reagent (Invitrogen) or PureLink™ RNA Mini Kit (Ambion) and treated with RNase-free DNase I (Roche) to remove contaminated genomic DNA. polyA$^-$ and polyA$^+$ RNAs were isolated from total RNA using the Poly(A)Purist™ MAG Kit (Ambion) following manufacturer's protocol. Isolation of RNA from subcellular fractions was performed as previously described [8] with modifications. In brief, cells were lysed in cytosolic lysis solution (10 mM HEPES pH 7.9, 1.5 mM $MgCl_2$, 10 mM KCl, 0.5 % NP40, 1 mM DTT plus protease and RNase inhibitors) for 10 minutes on ice. After centrifugation, the supernatant was collected as the cytoplasmic fraction for cytosolic RNA isolation. After washing in cytosolic lysis solution, the nuclear pellet was used for nuclear RNA isolation. To collect nucleoplasm and chromatin fractions, the nuclear pellet was further lysed with nuclear lysis solution (20 mM HEPES pH 7.9, 1.5 mM

MgCl2, 450 nM NaCl, 0.2 mM EDTA, 25% glycerol, 1 mM DTT, plus protease and RNase inhibitors). After centrifugation, the nuclear-soluble fraction (nucleoplasm) was collected as supernatant and the chromatin-associated fraction was collected as the pellet. RNAs from collected fractions were extracted with Trizol reagent and treated with RNase-free DNase I (Roche).

For RT-PCR, RNA was reverse-transcribed by using the SuperScript® III Reverse Transcriptase (Invitrogen). Red Taq Pro Complete (Denville Scientific) was used to amplify designated amplicons. For qPCR assays, cDNA was generated by the QuantiTect Rev. Transcription Kit (Qiagen) which also includes additional DNA contamination removal. iQ SYBR Green Supermix (Biorad) was used for PCR quantitation in a RotorGene cycler (Corbett). Relative quantification was performed using the ddCt method [9]. To calculate *LOUP* transcript numbers per cell, *LOUP* DNA fragments amplified by RT-PCR from HL-60 cDNA were cloned into the pSCAmpKan plasmid (Agilent). *LOUP* RNA fragments were *in vitro*-transcribed by using a MAXIscript™ Transcription Kit (Ambion). The RNA fragments were used to generate a standard curve for absolute quantification in qRT-PCR assays. Primers and probes used for all PCR assays are provided in Table S3.

**Fluorescence-activated cell sorting and analysis**

Cell populations were isolated for RNA extraction as previously described [10]. Briefly, mononuclear cells were isolated from bone marrow, spleen and peripheral blood after lysing red blood cells with ACK lysis buffer [11]. Single cell suspensions were stained with fluorochrome-conjugated antibodies (Biolegend and eBioscience) and FACS-sorted based on the following markers. LT-HSC: Lin$^-$c-Kit$^+$Sca-1$^+$CD150$^+$CD48$^-$; ST-HSC: Lin$^-$c-Kit$^+$Sca-1$^+$CD150$^-$CD48$^+$; LMPP: Lin$^-$c-Kit$^+$Sca-1$^+$CD34$^+$Flt3$^+$; MEP: Lin$^-$c-Kit$^+$Sca-1$^-$CD34$^-$CD16/32$^-$; CMP: Lin$^-$c-Kit$^+$Sca-1$^-$CD34$^+$CD16/32$^-$; GMP: Lin$^-$c-Kit$^+$Sca-1$^-$CD34$^+$CD16/32$^+$; Myeloid: Mac1$^+$Gr1$^+$. Myeloid surface marker staining and FACS analysis were performed following previously described procedures [12]. Cells were stained with PACBLUE-CD11b

(BioLegend). Stained cells were analyzed using LSRII flow cytometer (BD Biosciences) and FlowJo software (Tree Star)

**Transcript mapping by P5-linker ligation and 3' RACE**

The 5' end of the *LOUP* transcript was identified using the P5-linker ligation method as described previously [13]. Briefly, single-stranded cDNAs were generated from HL-60 polyA$^+$ RNA by using SuperScript III reverse transcriptase (Life Technologies) with *LOUP*-specific nested primer #1. Double-strand cDNAs were then synthesized from single-stranded cDNA using a SuperScript™ Double-Stranded cDNA Synthesis Kit (Life Technologies) and blunt-ended by NEBNext End Repair Enzym Module (New England Biolabs). After purification, these cDNAs were ligated with the P5-splinkerette adapter and purified. All purification steps were done by using QIAquick PCR Purification Kit (QIAGEN). Ligated products were then purified and used as templates for PCR using a P5 primer and *LOUP*-specific nested primers #1 and #2 with Phusion Hot Start DNA polymerase (Finnzymes). P5-linker ligation products were gel purified using a QIAgen Gel Extraction Kit (QIAGEN), sub-cloned into the pSCAmpKan vector, and transformed into competent bacteria using a StrataClone Blunt PCR Cloning Kit (Agilent). The 3'RACE assay was performed using a 2$^{nd}$ Generation 5'/3' RACE Kit (Roche) according to manufacturer's instructions. In brief, cDNA was generated from HL-60 polyA$^+$ RNA using oligo(dT)-anchor primer mix. Overlapping RACE products were then amplified from cDNA using an anchor primer and *LOUP*-specific primers. RACE products were sub-cloned into the pSCAmpKan vector and transformed into competent bacteria using a StrataClone Cloning Kit (Agilent). Plasmids containing P5-linker and RACE products were purified from bacteria, sequenced, and assembled. Primer information is in Table S3.

**Northern blotting**

10 ug polyA$^-$ and polyA$^+$ RNAs were dissolved and heat denatured in sample buffer containing formamide, MOPS and formaldehyde. Denatured RNAs were separated on a 1% denaturing agarose gel containing formaldehyde, MOPS and EtBr, before being transferred to Brightstar-plus positively charged nylon membrane (Life Technologies). The *LOUP* probe was PCR amplified with primers described in Table S3. The PCR product was sub-cloned into cloned into the pSCAmpKan vector using a StrataClone PCR Cloning Kit (Agilent). The probe sequence was verified by Sanger sequencing. The probe was released from the vector by restriction enzyme digestion and gene purification. The *LOUP* probe was radiolabeled using the Random Primed DNA Labeling Kit (Roche). Northern blot analysis was performed with ExpressHyb™ Hybridization Solution (Clontech) following the manufacturer's protocol.

**Quantitative Chromosome Conformation Capture (3C-qPCR)**

3C-qPCR experiments were performed by adapting described methods [14-16]. Briefly, 1x$10^6$ cells were crosslinked using 1% formaldehyde in PBS at room temperature for 10 minutes. The crosslinking reaction was stopped by adding 0.125 M Glycine and incubated for 5 minutes at room temperature followed by 15 minutes on ice. Crosslinked cells were then washed with ice-cold PBS and lysed in 3C lysis buffer (10 mM Tris-HCl, pH 8.0; 10 mM NaCl; Igepal CA-630 0.2% (vol/vol); 1X protease inhibitor cocktail (Sigma)) with 15 Dounce homogenizer strokes. After centrifugation, nuclear pellets were washed in 1x restriction enzyme buffer before being lysed with 0.1% SDS in 1x restriction enzyme buffer at 65$^0$C for 10 minutes. After incubation, the chromatin solution was supplemented with 1% Triton X-100 and digested by ApoI restriction enzyme (New England Biolabs) at 37$^0$C overnight with rotation. The following day, 1.5% SDS was added to the reaction and enzyme activity was inhibited by incubating at 65$^0$C for 30 minutes. Nearby DNA ends of digested chromatin were joined by T4-ligase (New England Biolabs) at 16$^0$C for 2 h. Bound proteins, including histones, were removed by proteinase K digestion at 65$^0$C overnight. The DNA library was extracted by phenol/chloroform using phase-lock gel tubes (5PRIME) and ethanol precipitation. RNA was removed by incubating 3C libraries with RNase A (Lucigen) at 37$^0$C for 15 minutes. TaqMan real-time PCR quantifications of ligation products were performed, using primers and probes as documented in Table S3.

**Chromatin Isolation by RNA Purification (ChIRP)**

ChIRP assays were performed as previously described [17,18] with additional modifications. Briefly, to preserve RNA-chromatin interactions, cells were first crosslinked with 2 mM EGS at room temperature for 45 minutes. After washing cells with ice-cold PBS, cells were further crosslinked with 3% paraformaldehyde for 15 minutes at room temperature. The crosslinking reaction was quenched with 0.125 M glycine for 5 minutes at room temperature. Crosslinked cells were washed in ice-cold PBS and lysed in sonication buffer (20 mM Tris pH 8, 150 mM NaCl, 0.1% SDS, 1% Triton-X, 2 mM EDTA, 1 mM PMSF) supplemented with cOmplete™, Mini Protease Inhibitor Cocktail (Sigma-Aldrich) and SUPERase In RNase Inhibitor (Invitrogen). After sonication and centrifugation, the supernatant containing sheared chromatin was collected and incubated with biotinylated anti-sense DNA tiling probes in hybridization buffer (750 mM NaCl, 1% Triton, 0.1% SDS, 50 mM Tris-Cl pH 7.0, 1 mM EDTA, 15% formamide, 1 mM PMSF) supplemented with cOmplete™, Mini Protease Inhibitor Cocktail and SUPERase In RNase Inhibitor. Hybridized chromatin fragments were captured using Dynabeads™ MyOne™ Streptavidin C1 (Invitrogen). Captured chromatin fragments was either used for extracting chromatin-bound RNA by Trizol reagent or for DNA isolation. Chromatin-bound *LOUP* was quantitated from chromatin-bound RNA by qRT-PCR. Enrichment of the URE and the PrPr were evaluated by qPCR. Probes used in the ChIRP assay were designed using the online probe designer at singlemoleculefish.com and are listed in Table S3.

**DNA pull-down assay (DNAP)**

DNAP was performed as described previously with minor modifications [19]. Briefly, the nuclear extract was pre-cleared with Dynabeads™ MyOne™ Streptavidin C1 for 30 minutes at 4°C then incubated overnight with biotinylated oligonucleotides in binding buffer (10 mM HEPES pH 7.9; 100 mM KCl, 5 mM MgCl2, 1 mM EDTA, 10% glycerol, 1 mM DTT, 0.5% NP-40, 1 mM DTT) supplemented with 1x protease inhibitor cocktail (Sigma-Aldrich). Beads were washed with binding buffer then added to the binding reaction. After 1-hour incubation, beads were washed five times with binding buffer. DNA-bound proteins were eluted from beads and subjected to SDS-PAGE and immunoblotting.

**RNA pull-down assay (RNAP) and RNA-Protein interaction prediction**

RNAP were performed essentially as described previously [20] with few modifications. Briefly, biotinylated RNA was *in vitro*-transcribed using the MAXIscript™ Transcription Kit (Ambion). The DNA template was removed by DNAseI treatment. Transcribed RNA was purified using a RNeasy Mini Kit (QIAGEN). Purified RNA was denatured by heating to 90°C for 2 minutes following incubation on ice for 2 minutes in RNA structure buffer (10 mM Tris pH 7, 0.1 M KCl, 10 mM MgCl2). Denatured RNA was then shifted to room temperature for 20 minutes to form proper secondary structure. Nuclear extracts were treated with RNase-free DNase I (Roche) to remove genomic DNA and pre-cleared with Dynabeads™ MyOne™ Streptavidin C1 or Streptavidin agarose beads (Invitrogen) in binding buffer I (150 mM KCl, 25 mM Tris pH 7.4, 0.5 mM DTT, 0.5% NP40, 1 mM PMSF) supplemented with cOmplete™, Mini Protease Inhibitor Cocktail, and SUPERase In RNase Inhibitor. Pre-cleared extracts were then incubated with biotinylated RNAs in binding buffer I for 1 hour. Beads were washed with binding buffer I then added to the binding reaction. After 1-hour incubation, beads were washed five times with binding buffer I. RNA-bound proteins were eluted from beads and subjected to SDS-PAGE and immunoblotting. For recombinant proteins, binding buffer II (50 mM Tris-Cl 7.9, 10% Glycerol, 100 mM KCl, 5 mM MgCl2, 10 mM β-ME, and 0.1% NP-40) was used.

*In silico* prediction of RNA-Protein interactions were performed using the catRAPID Fragments algorithm in which protein-RNA interaction propensities were predicted  based on calculation of secondary structure, hydrogen bonding, and van der Waals contributions [21]

**RNA Immunoprecipitation sequencing and qPCR (RIP-seq and RIP-qPCR)**

RIP was performed following a protocol reported by Hendrickson et al [22] with modifications. Briefly, cells were crosslinked in 0.1% formaldehyde at room temperature for 10 minutes. The crosslinking reaction was quenched for 5 min at room temperature with 0.125 M glycine. Crosslinked cells were washed with ice-cold PBS. Cell pellet was lysed in RIPA lysis buffer (50 mM Tris (pH 8), 150 mM KCl, 0.1 % SDS, 1 % Triton-X, 5 mM EDTA, 0.5 % sodium deoxycholate, 0.5 mM DTT) supplemented with protease inhibitor cocktail (Thermo Scientific) and 100 U/ml RNaseOUT™ (Invitrogen). After sonication, cell lysate was pre-cleared by incubating with Dynabeads® Protein G (Invitrogen). Beads were then captured and removed using a magnet. Pre-cleared lysate was incubated with anti-RUNX1 antibody or IgG (Abcam) at 4°C for 2 hours before adding 50 µl of Dynabeads® Protein G to capture antibodies. After washing, beads were kept at -20$^0$C or proceeded to incubation with reverse-crosslinking buffer (3× PBS (without Mg or Ca), 6 % N-lauroyl sarcosine, 30 mM EDTA, 15 mM DTT) supplemented with Proteinase K (Ambion) and RNaseOUT together with the input sample. Captured RNAs were extracted with Trizol reagent. Contaminated DNA was removed from extracted RNA by DNAseI from RNase-Free DNase Set (QIAGEN) then ribosomal RNA was removed using the Ribo-Zero™ Magnetic Gold Kit (Epicentre). RNA was further purified using RNeasy MinElute Cleanup Kit (QIAGEN). RNA quality was determined using the RNA 6000 Pico Kit on a Bioanalyzer (Agilent). Purified RNA was used for qRT-PCR as described elsewhere and cDNA library construction using the Truseq stranded total RNA library prep kit (Illumina) according to the manufacturer’s protocol. The libraries were pooled together and subjected to pair-end sequencing on a Nextseq500 (Illumina) to achieve 2×40 bp reads.

**Chromatin Immunoprecipitation and qPCR (ChIP-qPCR)**

ChIP was performed as previously described [23]. Briefly, 2x10$^6$ U937 cells were crosslinked with 1% formaldehyde (formaldehyde solution, freshly made: 50 mM HEPES-KOH; 100 mM NaCl; 1 mM EDTA; 0.5 mM EGTA; 11% formaldehyde) for 10 minutes at room temperature. The crosslinking reaction was stopped by incubating with 0.125 M glycine for 5 minutes at room temperature. Crosslinked cells were washed twice with ice-cold PBS (freshly supplemented with 1 mM PMSF). Cell pellet was lysed for 10 minutes on ice and chromatin was fragmented by sonication (25 cycles, 30 seconds on, 60 seconds off, high power, Bioruptor). The chromatin solution was incubated with 10 µg antibody overnight at 4°C. Protein A magnetic beads (NEB) were used to capture antibody-bound chromatin. After washing, chromatin was reverse-crosslinked and treated with proteinase K overnight at 65°C. Beads were then removed using a magnet and the chromatin solution was treated with RNase treatment (Epicentre) for 30 minutes at 37°C. ChIP DNA was extracted with Phenol:Chloroform:Isoamyl Alcohol 25:24:1, pH:8 (Sigma-Aldrich) and then precipitated with an equal volume of isopropanol in the presence of glycogen. The DNA pellet was dissolved in 30 µl of TE buffer for qPCR analyses. Fold enrichment was calculated using the formula $2^{(-\Delta\Delta Ct(ChIP/IgG))}$. Primer sets used for ChIP-qPCR are listed in Table S3.

**RIP-seq and ChIP-seq data analyses**

RIP-seq samples were demutliplexed. Reads were deduplicated by Clumpify from the BBtools suite, sourceforge.net/projects/bbmap/) with the parameters “dedupe spany addcount”. Adaptor quality trimming and filtering was performed by BBDuck from the BBtools suite with the parameters “ktrim=l hdist=2”. Low quality reads/bases were removed by Trimmomatic [24] with the parameters: “LEADING:28 SLIDINGWINDOW:4:26 TRAILING:28 MINLEN:20”. The processed reads were then aligned to Human genome build 38 (hg38) by the STAR aligner [25] with the parameters “--outFilterScoreMinOverLread 0.05 --outFilterMatchNminOverLread 0.05 --outFilterMultimapNmax 30 --outSAMprimaryFlag AllBestScore”. Coverage maps were generated using bamCoverage (part of the deepTools suite [26] with default parameters. Peak calling was performed using HOMER (v4.10) [27]. RUNX1 peaks with at least ten-fold enrichment over the surrounding 10 kb region were selected for annotation using HOMER. Peaks were assigned to a gene locus by satisfying at least one of the following location criteria: a nearest transcription start site, on a promoter, or on a transcript body. Ensemble 97 human gene CRCh38.p12 was used to retrieved gene annotation information through Biomart in Ensembl [28]. For ChIP-seq and Dnase-seq data, raw reads were downloaded from GEO (RUNX1 ChIP-seq in THP-1 cells: GSM2108052; RUNX1-ETO and H3K9Ac ChIP-seq, and Dnase-seq in Kasumi-1: GSE29222). Read quality were evaluated by FastQC [29]. Where necessary, reads with low-quality were trimmed by trim_galore [30]. Genome alignment, coverage maps, and peak calling were performed using software packages as above. BigWig files were uploaded and viewed via the UCSC genome browser.

The following gene tracks were from published data deposited in GEO and were processed via the Cistrome pipeline [31]. H3K27ac overlay track includes monocytes (GSM2679933), THP-1 (GSM2544236), and HL-60 (GSM2836486). The H3K4me1 overlay track includes monocyte (GSM1435532), HL-60 (GSM2836484) and THP-1 (GSM3514951). H3K4me3 overlay track includes monocytes (GSM1435535), HL-60 (GSM945222), and THP-1 (GSM2108047). The DNAse-seq overlay track includes monocytes (GSM701541) and HL-60 (GSM736595). RUNX1 ChIP-seq tracks include $CD34^+$ cells from healthy donors (GSM1097884), an AML patient with FLT3-ITD and no other defined mutations (GSM1581788), and an AML patient with non-t(8;21) (GSM722708). The CAGE track (reverse strand and max counts) was imported from the FANTOM5 project [32].

**RNA sequencing data analysis (RNA-seq)**

Raw sequencing reads (FASTQ files) of the Human Body Map data set were downloaded from AEArrayExpress (E-MTAB-513). Read quality was assessed by FastQC [29]. Reads with low-quality were trimmed by trim_galore [30]. The *LOUP* transcript was integrated into the Ensembl human cDNA catalog GRCh38 and transcript levels were quantified against this catalog using the Salmon software [33]. AML RNA-seq were downloaded from TCGA and transcript counts were determined. For RNA-seq track visualization, the following RNA-seq raw data were downloaded from GEO: THP-1 (GSM1843218), HL-60 (GSM1843216), $CD34^+$ HSPC (GSM1843222), Monocyte (GSM1843224), and Jurkat (GSM2260195). BigWig files were generated using packages as described in ChIP-seq and RIP-seq analyses and viewed via the UCSC genome browser.

**Single-cell RNA-seq (scRNA-seq) data analyses**

Raw fastq files data of mononuclear cells isolated from peripheral blood and bone marrow were obtained from the 10x Genomics public datasets repository (https://www.10xgenomics.com/resources/datasets/) and pooled together. Transcripts were mapped to the human transcriptome using Cell Ranger (10x Genomics) with a custom hg38 gtf containing the *LOUP* transcript details. Subsequent analyses were performed in R (v3.6.2) using the previously published Bioconductor workflow with minor modifications [34]. Filtering criteria were as below. First, cells with library sizes more than three median absolute deviations (MADs) below the median library or four MAD's above the median library size were filtered out. Second, cells with a total number of expressed genes (>= 1 read) more than three MADs below the median total number of expressed genes or four MAD's above the median total number of expressed genes were filtered out. Third, cells with a total percentage of expressed genes originating from mitochondrial DNA more than eight MADs above the median were filtered out. A doublet score was then computed to estimate the percentage of barcodes for two or more cells as previously described [35]. Cells with a doublet score of 0.99 were excluded. Expression of each cell was normalized by a size factor approach as previously described [36] resulting in $log_2$(normalize_expression) values. Principle component and t-Distributed Stochastic Neighbor Embedding (tSNE) analyses revealed no significant batch effects to be regressed out for the samples. To account for dropouts which are being more frequent for genes with lower expression magnitude in scRNA-seq [37], cells with undetectable *LOUP* and *PU.1* transcripts were referred as $LOUP^{low}/PU.1^{low}$ and cells with detectable *LOUP* and *PU.1* transcripts were referred as $LOUP^{high}/PU.1^{high}$. Expression data visualization was performed using SPRING software [38]. Briefly, a graph of cells connected to their nearest neighbors in gene expression space was determined. This was then projected into two dimensions using a force-directed graph layout. Identity of each cell was inferred using the Blueprint-Encode annotation which includes normalized expression values of 259 bulk RNA-seq samples generated from pure and defined cell populations [39,40]. This annotation was integrated in the SingleR R package [41]. Annotated cells were grouped into major definitive cell lineages as described in the text.

Gene Ontology (GO) analysis was performed using the Database for Annotation, Visualization and Integrated Discovery functional annotation tool (http://david.abcc.ncifcrf.gov). Significance of over-represented Gene Ontology biological processes was examined based on $-\log_{10}$ of corrected *p*-values from Bonferroni-corrected modified Fisher's exact test [42]. A list of enriched genes in *LOUP*$^{high}$/*PU.1*$^{high}$ group *vs.* *LOUP*$^{low}$/*PU.1*$^{low}$ group was generated using SPRING software [38]. Upregulated genes (Z-score >1) was used for GO analysis.

**Prediction of coding potential with PhyloCSF**

The cross-species multiple sequence comparisons result of 46 species (i.e., multiz100way) was downloaded from the UCSC genome browser (https://genome.ucsc.edu). Guided by the GENCODE gene annotation (ver. 28), the alignment of the longest isoform of each gene was extracted from alignments of cross-species multiple sequence comparisons. The alignment was analyzed by PhyloCSF [43] with 58mammals mode. All possible coding reading frames on the same strand were scanned. The maximal score was used.

### SUPPLEMENTAL FIGURES

**Figure S1. Identification of gene loci exhibiting concurrent RUNX1-RNA and -DNA interactions, Related to Figure 1**

(A) Workflow of RUNX1-RIP procedure. Ab: antibody.

(B) Immunoblot detection of RUNX1 and actin proteins immunoprecipitated from THP-1 cell lysate using anti-RUNX1 antibody and Rabbit IgG isotype control.

(C) Bioanalyzer analysis of RNAs captured by anti-RUNX1 antibody and IgG control plus input RNAs.

(D) Analysis flowchart of RUNX1 RIP-seq and ChIP-seq analyses.

(E and F) Pie charts showing distribution of RUNX1 RIP-seq peaks and RUNX1 ChIP-seq peaks at different genomic locations.

(G) Examples of the myeloid gene loci having both RUNX1 RIP peaks and RUNX1 ChIP-seq peaks from THP-1 cells.

**Figure S2. Transcript map and molecular features of *LOUP*, Related to Figure 2**

(A) RT-PCR confirmation of exon-exon junction. Upper panel: Schematics of the PCR amplicon and primer locations. Lower panels: DNA sequencing of PCR products from human (HL-60) and murine (RAW264.7) cells.

(B) Workflow of 5' end mapping by the P5-linker ligation method.

(C) P5-linker ligation assay determining the 5' end of *LOUP* transcript. Upper panel: DNA sequencing analysis showing locations of the P5-primer, P5-splinkerette, and transcription start site (TSS). Lower panel: Schematic diagram of the *PU.1* locus. Shown are the URE element with the two homology regions H1 and H2.

(D) Schematic diagram showing the relative genomic location of *LOUP* and two neighbor genes: *PU.1* and *SLC39A13* (top), the splicing pattern of *LOUP* (middle), and resultant transcripts (bottom). E1: Exon 1, E2: Exon 2. E2a and E2b are exons derived from an additional splicing event within Exon 2. Exon boundaries were mapped by 3'RACE and RT-PCR.

(E) PhyloCSF analysis of *LOUP* and other known coding and noncoding genes. Shown are coding potential scores.

(F) qRT-PCR analysis of *Loup* RNA in subcellular fractions isolated from RAW264.7 cells. Fraction enrichment controls include *Malat1* (chromatin) and *Rps18* (cytoplasm) [44].

(G) qRT-PCR analysis of fraction enrichment controls including *MALAT1* (polyA$^{+}$) and *RPPH1* (polyA$^{-}$) (right panel).

(H) Measurement of transcript numbers per HL-60 cell. Upper panel: Schematic diagram of amplified amplicons showing primer locations for non-spliced *LOUP* (FW2-RV) and spliced *LOUP* (FW1-RV). Lower panels: qRT-PCR with RNA standard curve for spliced and non-spliced forms.

(I) Left panel: qRT-PCR analysis of *LOUP* forms in the nucleus. Right panel: Fraction enrichment controls include *MALAT1* (nucleoplasm) and *RPS18* (cytoplasm).

Error bars indicate SD (n=3).

**Figure S3. Gene expression profiles in normal tissues and cell lineages, Related to Figure 3**

(A and B) Transcript profiles of *SLC39A13* and *RUNX1* in human tissues. Shown are transcript counts from the Illumina Body Map dataset. Error bars indicate SD (n=2).

(C) SRING plot analysis of the 10x Genomic scRNA-seq dataset showing color-coded definitive blood lineages using the Blueprint-Encode annotation [41]. Annotated cells in sub-populations were grouped into major cell populations (see methods for details).

(D, E and F) Transcript profiles of *LOUP* RNA, and *PU.1* and *RUNX1* mRNAs in blood cell lineages of the 10x Genomic scRNA-seq dataset. Each dot on the graph represents an individual cell.

(G) GO analysis for enrichment of biological processes using a list of genes upregulated in *LOUP*$^{high}$/*PU.1*$^{high}$ cells as compared to *LOUP*$^{low}$/*PU.1*$^{high}$ cells. Top enrichment GO terms are shown.

**Figure S4. Effects of gain and loss of *LOUP* expression, Related to Figure 4**

(A) Schematic of strategy for *LOUP* depletion. Included is a FACS sorting scheme for isolation of cells expressing both mCherry (Cas9) and eGFP (sgRNAs).

(B and C) Inference of CRISPR edits (ICE) analyses for indel composition and frequency of CRISPR/Cas9 cell clones. Top panels: Trace file segments of amplified genomic regions surrounding sgRNA-binding sites (#D1 and #D2 *LOUP* sgRNAs) in the edited (upper panel) and the control (lower panel) samples. Dotted red underline: Protospacer adjacent motif (PAM) sequence. Solid black underline: guide sequences. Expected cut sites are denoted as vertical dotted lines. Bottom-left panel: Indel efficiency analysis. Bottom-right panel: Indel distribution analysis. Dashed lines indicate deletion length.

(D) Genomic PCR and Sanger sequencing confirmation of U937 cell clones with *LOUP* homozygous indels (L2a and L2b) and control (N1).

**Figure S5. *LOUP* interacts with RUNX1, Related to Figure 6**

(A) Immunoblot of RUNX1 and control proteins in nuclear and cytosol fractions from U937 cells.
(B) Nucleotide identity plot generated from alignment of *LOUP* to itself using discontinuous megablast algorithm from BLAST (http://blast.ncbi.nlm.nih.gov/). Boxed area depicts a repetitive region (RR) of 670 bp. The schematic diagram underneath illustrates mature *LOUP* lncRNA containing the RR region
(C) *In silico* prediction of RR-RUNX1 interaction by catRAPID Fragments algorithm. R1 and R2: two regions with high interaction scores.

### SUPPLEMENTAL TABLE LEGENDS

**Table S1. List of myeloid genes, Related to Figure 1.** 78 myeloid genes defined by their known roles in myeloid development or myeloid molecular markers.

**Table S2. List of enriched genes in *LOUP*$^{high}$/*PU.1*$^{high}$ cells, Related to Figure 3.**

**Table S3. Table of oligonucleotide information.** Primers, probes, and sgRNAs for various assays used in this article.

### QUANTITATION AND STATISTICAL ANALYSIS

In general, quantitation and statistical tests were performed using GraphPad Prism 8.0 software. Data are shown as mean ± SD. The paired two-tailed Student's t-test (otherwise specified in respective figure legends) was used to calculate statistical significance of differences between two experimental groups. $p \leq 0.05$ was considered statistically significant.

### DATA SETS

Data are available on the Gene Expression Omnibus database under GEO Series accession number GEO: GSE140459.

### CONTACT FOR REAGENT AND RESOURCE SHARING

Further information and request for resources and reagents should be directed to and will be fulfilled by the Lead Contacts, Daniel G. Tenen and Bon Q. Trinh

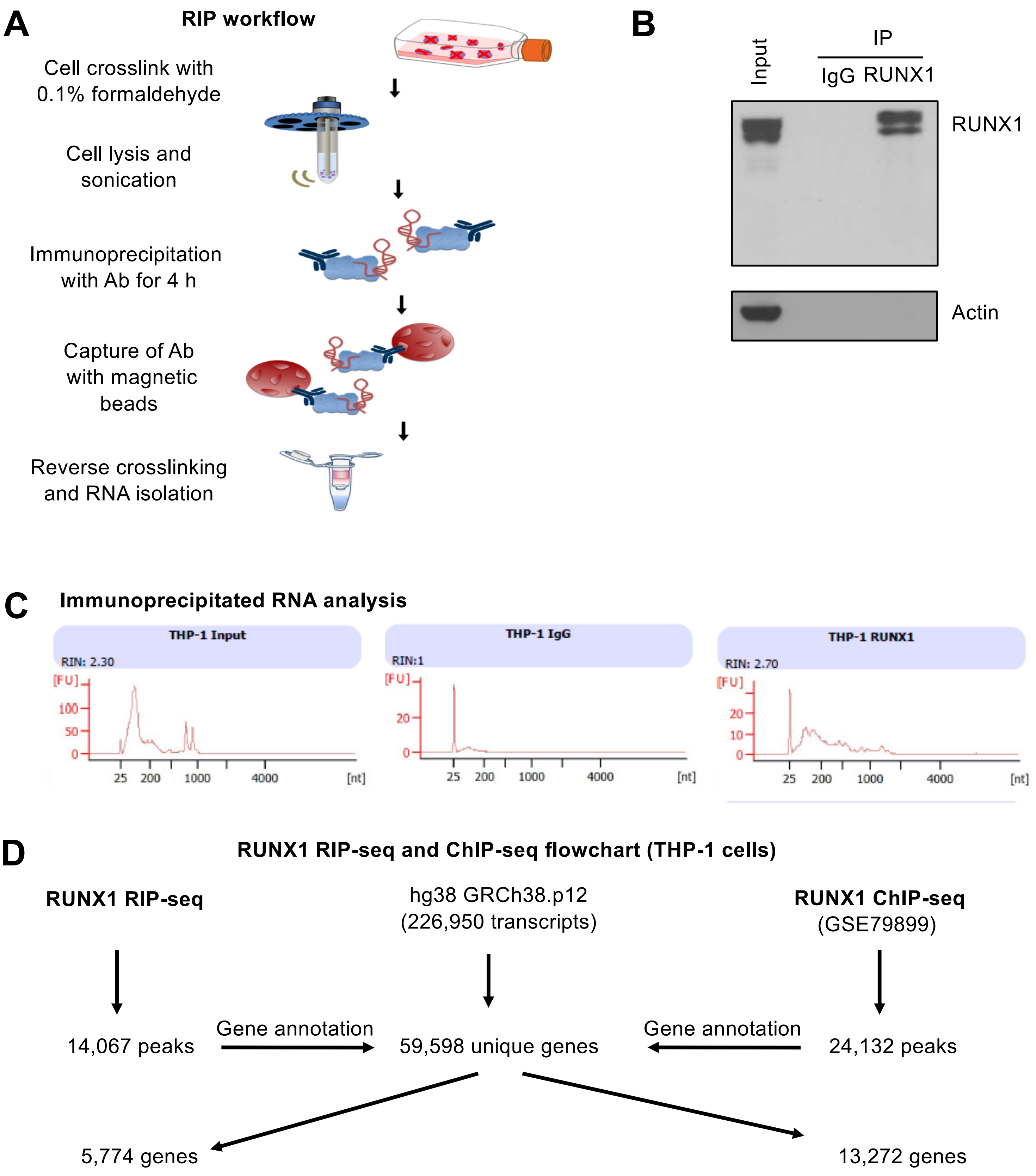
A
RIP workflow
Cell crosslink with
0.1% formaldehyde
Cell lysis and
sonication
Immunoprecipitation
with Ab for 4 h
Capture of Ab
with magnetic
beads
Reverse crosslinking
and RNA isolation
B
Input
IP
IgG RUNX1
RUNX1
Actin
C
Immunoprecipitated RNA analysis
THP-1 Input
RIN: 2.30
[FU]
100
50
0
25 200 1000 4000 [nt]
THP-1 IgG
RIN:1
[FU]
20
0
25 200 1000 4000 [nt]
THP-1 RUNX1
RIN: 2.70
[FU]
20
10
0
25 200 1000 4000 [nt]
D
RUNX1 RIP-seq and ChIP-seq flowchart (THP-1 cells)
RUNX1 RIP-seq
hg38 GRCh38.p12
(226,950 transcripts)
RUNX1 ChIP-seq
(GSE79899)
14,067 peaks
Gene annotation
59,598 unique genes
Gene annotation
24,132 peaks
5,774 genes
13,272 genes

**FigS1**

E

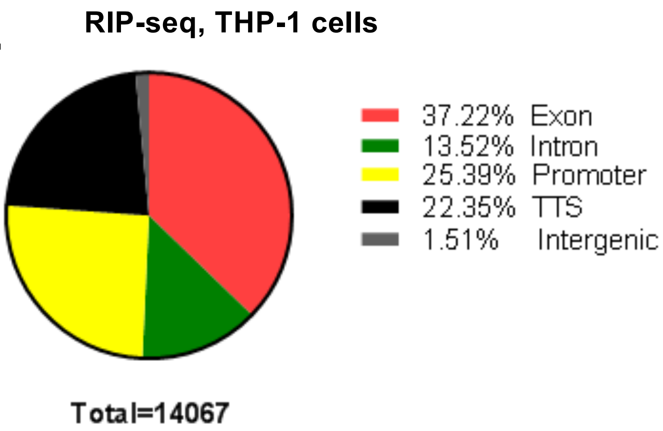
RIP-seq, THP-1 cells
37.22% Exon
13.52% Intron
25.39% Promoter
22.35% TTS
1.51% Intergenic
Total=14067

F

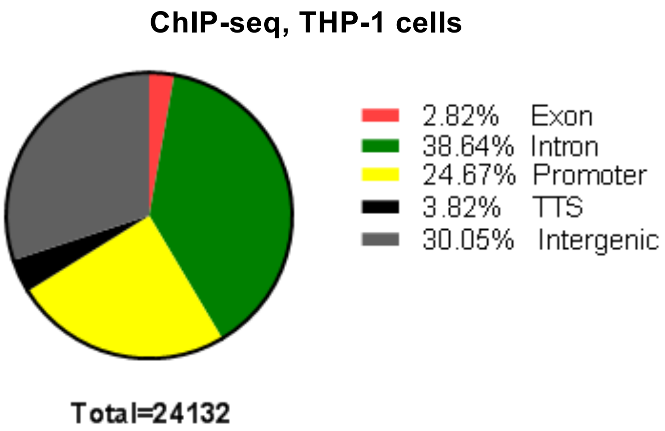
ChIP-seq, THP-1 cells
2.82% Exon
38.64% Intron
24.67% Promoter
3.82% TTS
30.05% Intergenic
Total=24132

G

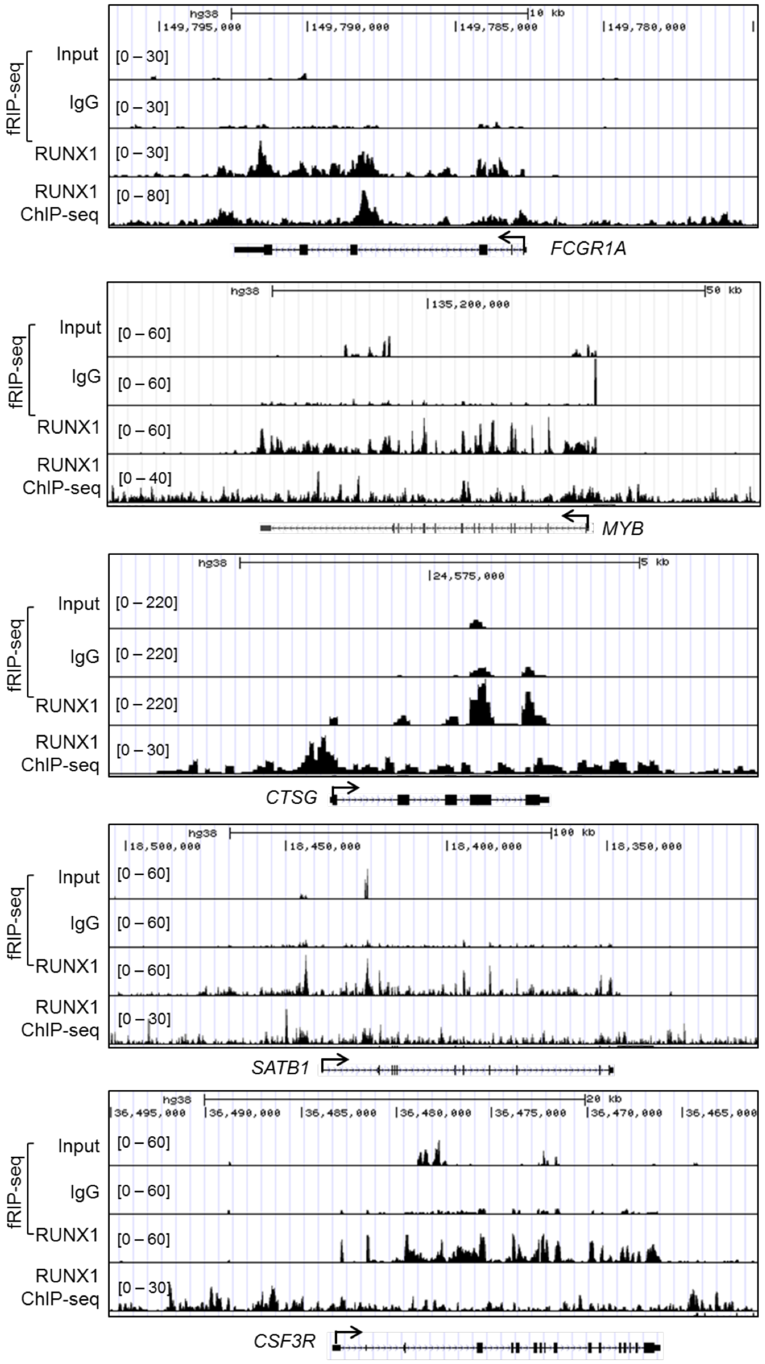
hg38
10 kb
149,795,000
149,790,000
149,785,000
149,780,000
fRIP-seq
Input
IgG
RUNX1
RUNX1
ChIP-seq
[0 – 30]
[0 – 30]
[0 – 30]
[0 – 80]
FCGR1A
hg38
50 kb
135,200,000
fRIP-seq
Input
IgG
RUNX1
RUNX1
ChIP-seq
[0 – 60]
[0 – 60]
[0 – 60]
[0 – 40]
MYB
hg38
5 kb
24,575,000
fRIP-seq
Input
IgG
RUNX1
RUNX1
ChIP-seq
[0 – 220]
[0 – 220]
[0 – 220]
[0 – 30]
CTSG
hg38
100 kb
18,500,000
18,450,000
18,400,000
18,350,000
fRIP-seq
Input
IgG
RUNX1
RUNX1
ChIP-seq
[0 – 60]
[0 – 60]
[0 – 60]
[0 – 30]
SATB1
hg38
20 kb
36,495,000
36,490,000
36,485,000
36,480,000
36,475,000
36,470,000
36,465,000
fRIP-seq
Input
IgG
RUNX1
RUNX1
ChIP-seq
[0 – 60]
[0 – 60]
[0 – 60]
[0 – 30]
CSF3R

**FigS1**

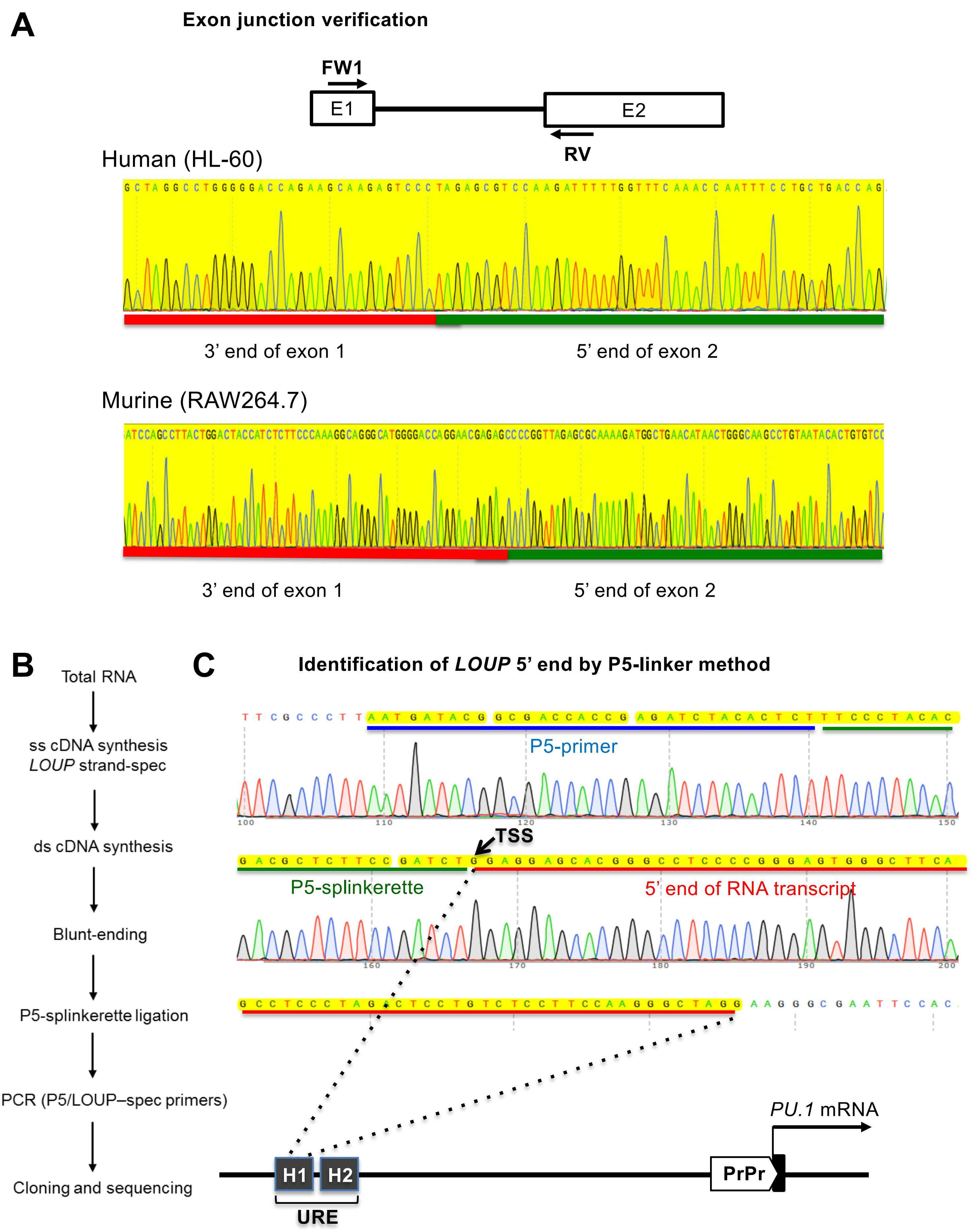
A
Exon junction verification
FW1
E1
E2
RV
Human (HL-60)
3' end of exon 1
5' end of exon 2
Murine (RAW264.7)
3' end of exon 1
5' end of exon 2
B
Total RNA
ss cDNA synthesis
LOUP strand-spec
ds cDNA synthesis
Blunt-ending
P5-splinkerette ligation
PCR (P5/LOUP–spec primers)
Cloning and sequencing
C
Identification of LOUP 5' end by P5-linker method
P5-primer
TSS
P5-splinkerette
5' end of RNA transcript
PU.1 mRNA
H1
H2
URE
PrPr

FigS2

D

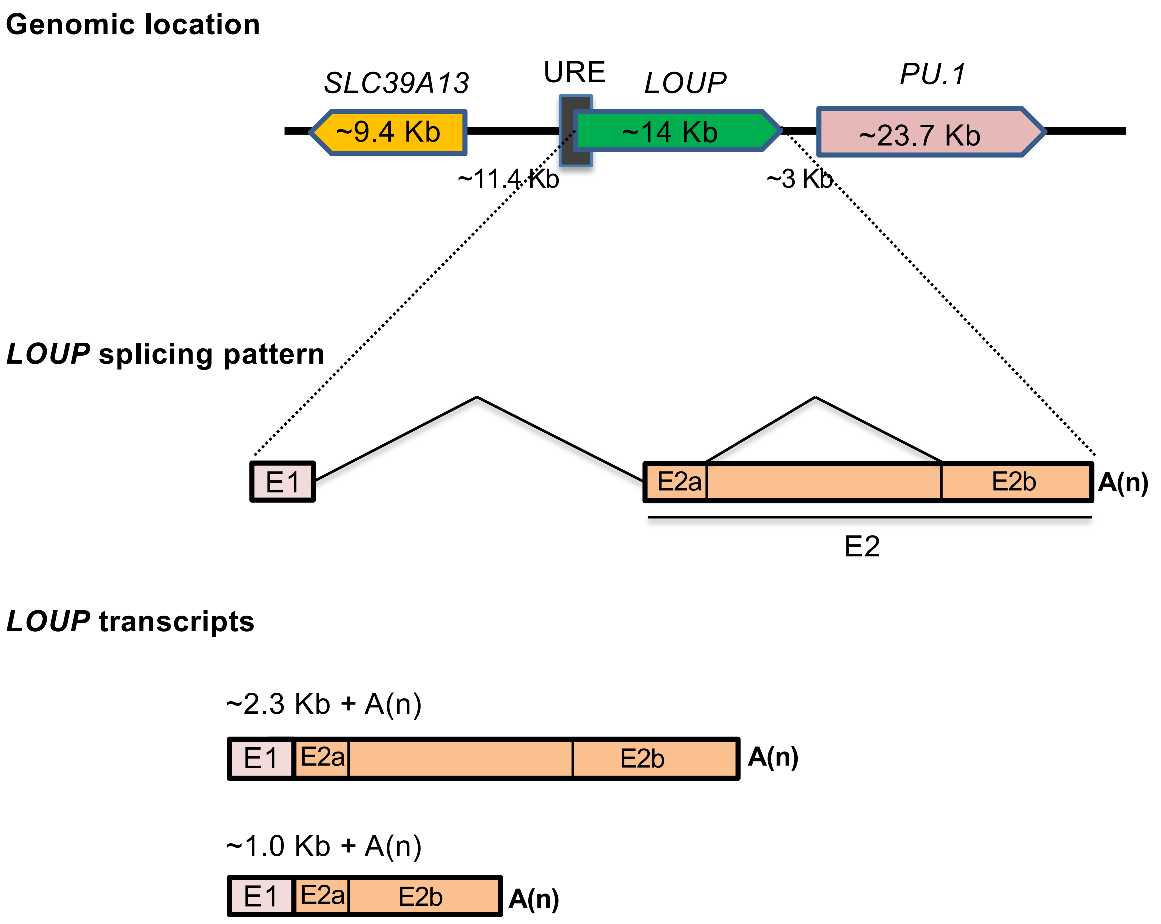
Genomic location
SLC39A13
URE
LOUP
PU.1
~9.4 Kb
~14 Kb
~23.7 Kb
~11.4 Kb
~3 Kb
LOUP splicing pattern
E1
E2a
E2b
A(n)
E2
LOUP transcripts
~2.3 Kb + A(n)
E1
E2a
E2b
A(n)
~1.0 Kb + A(n)
E1
E2a
E2b
A(n)

E

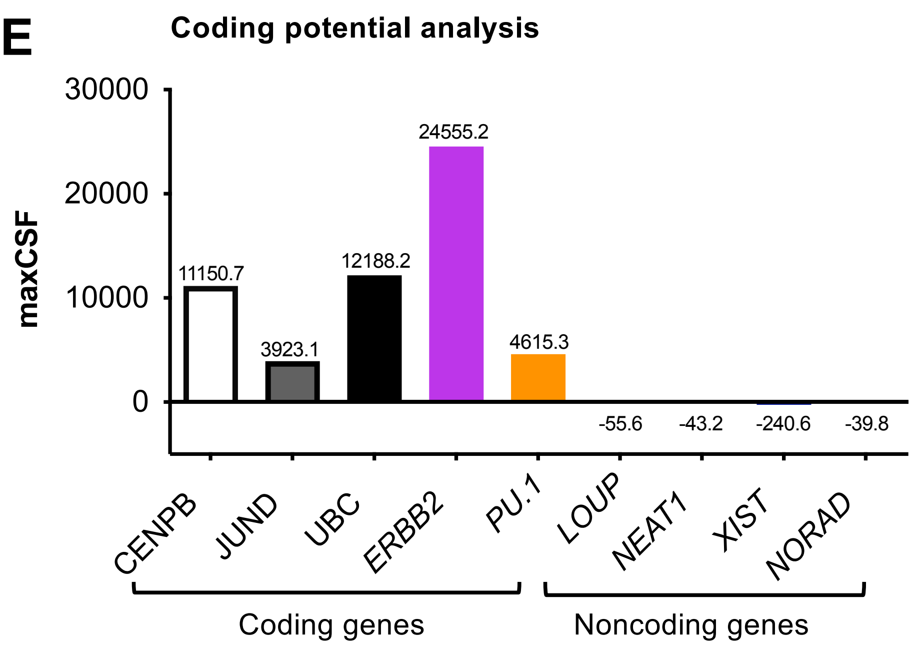
Coding potential analysis
maxCSF
30000
20000
10000
0
11150.7
3923.1
12188.2
24555.2
4615.3
-55.6
-43.2
-240.6
-39.8
CENPB
JUND
UBC
ERBB2
PU.1
LOUP
NEAT1
XIST
NORAD
Coding genes
Noncoding genes

FigS2

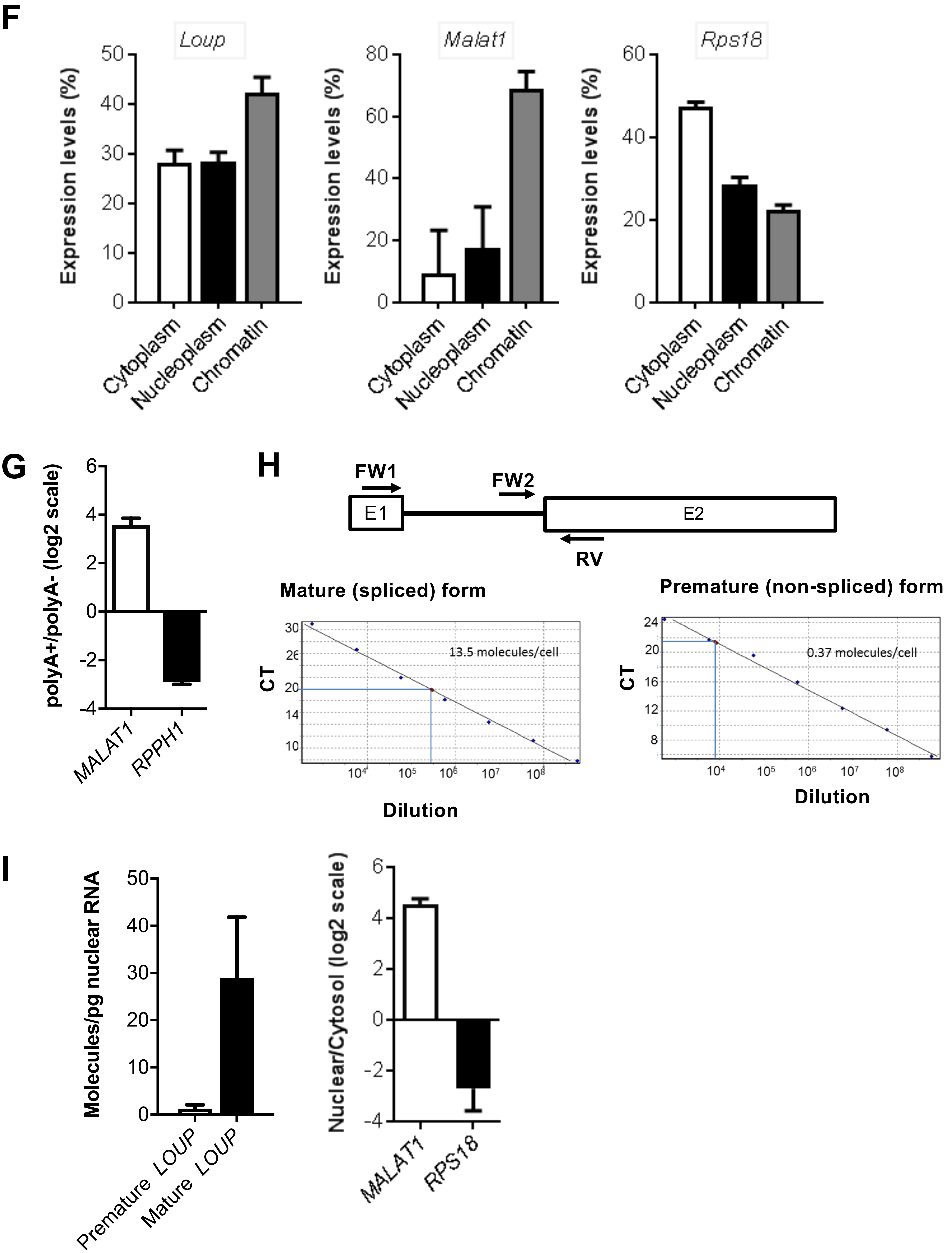
F
Loup
Malat1
Rps18
Expression levels (%)
Cytoplasm
Nucleoplasm
Chromatin
G
polyA+/polyA- (log2 scale)
MALAT1
RPPH1
H
FW1
FW2
E1
E2
RV
Mature (spliced) form
13.5 molecules/cell
CT
Dilution
Premature (non-spliced) form
0.37 molecules/cell
I
Molecules/pg nuclear RNA
Premature LOUP
Mature LOUP
Nuclear/Cytosol (log2 scale)
MALAT1
RPS18

FigS2

Bulk RNA-seq (Illumina Body Map)

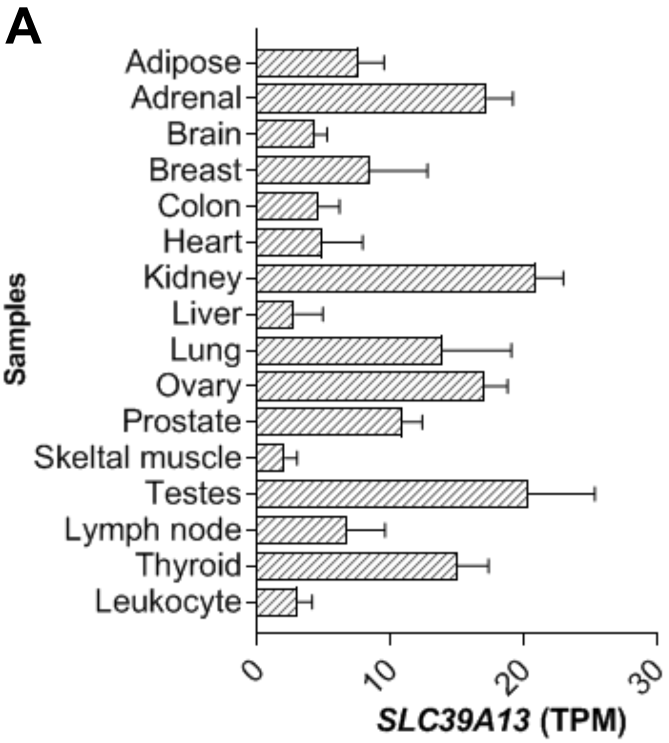


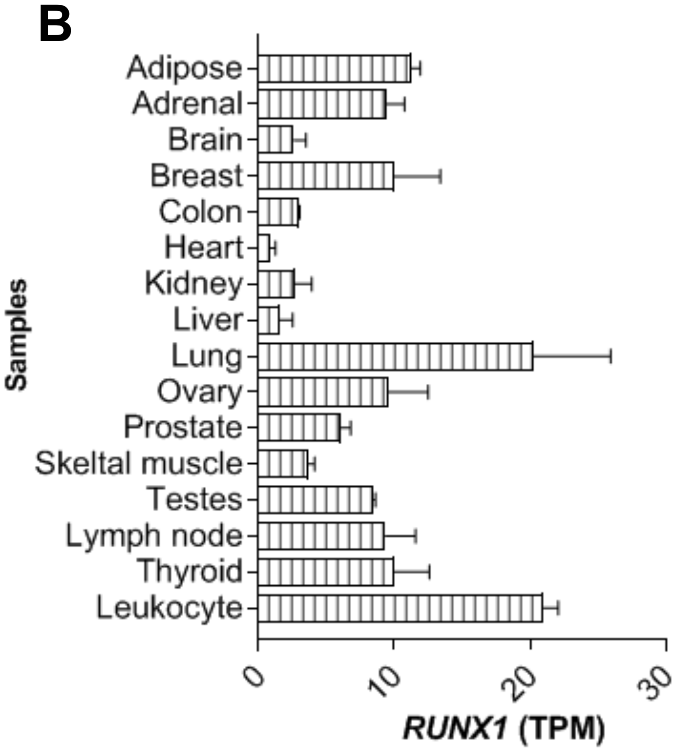


C

**Spring plot of human PMNC and BMNC (scRNA-seq, 10x Genomics)**

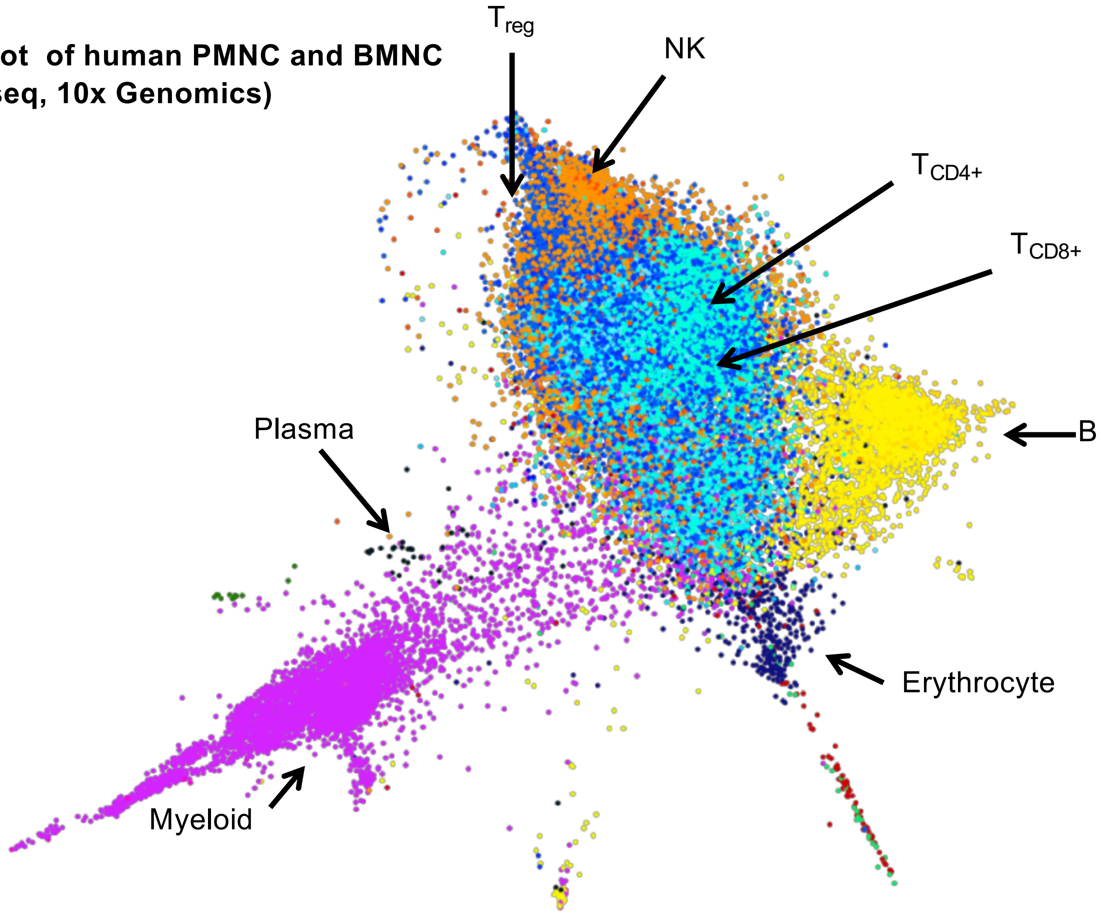


**FigS3**

scRNA-seq (10x Genomic)

D

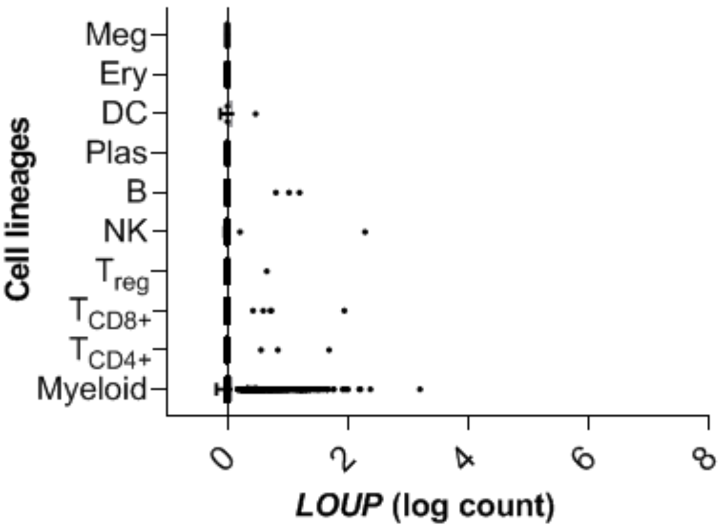


E

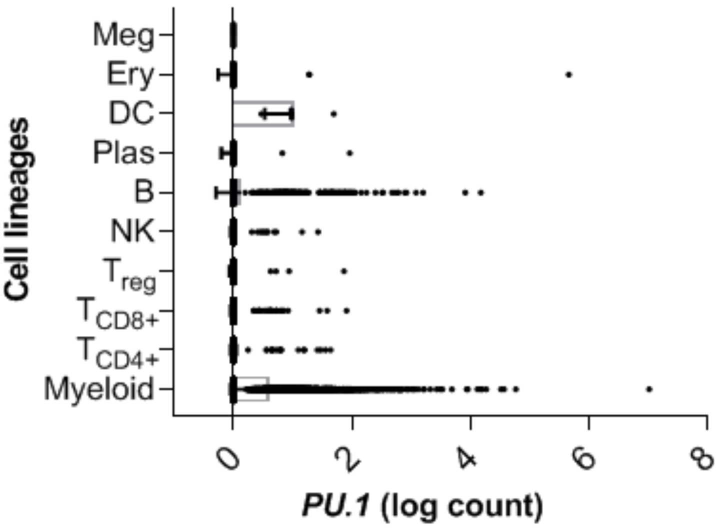


F

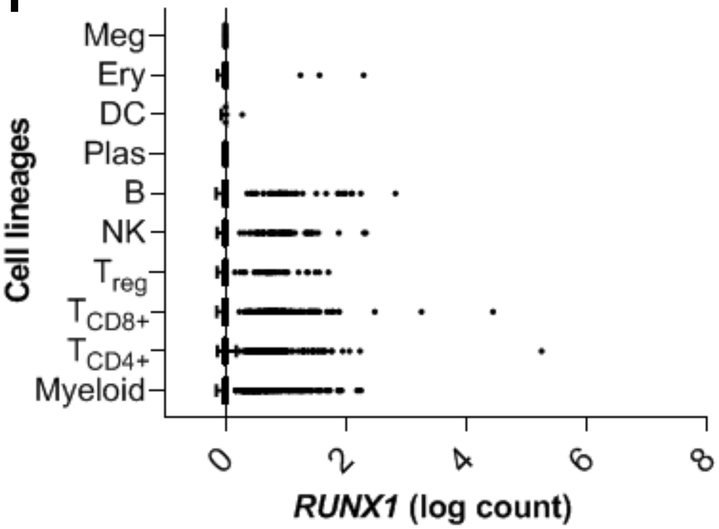


G

**GO analysis of genes enriched in *LOUP*$^{high}$/*PU.1*$^{high}$ cells (10x Genomic)**

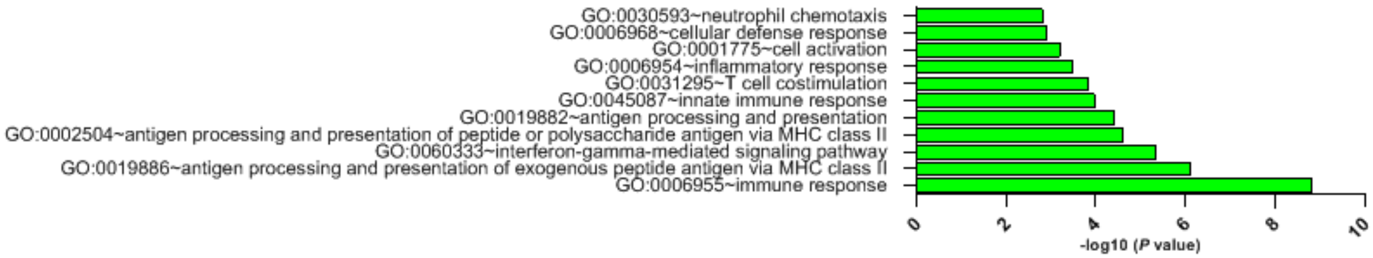


**FigS3**

**A** **Schematic strategy for *LOUP* depletion by CRISPR/Cas9**

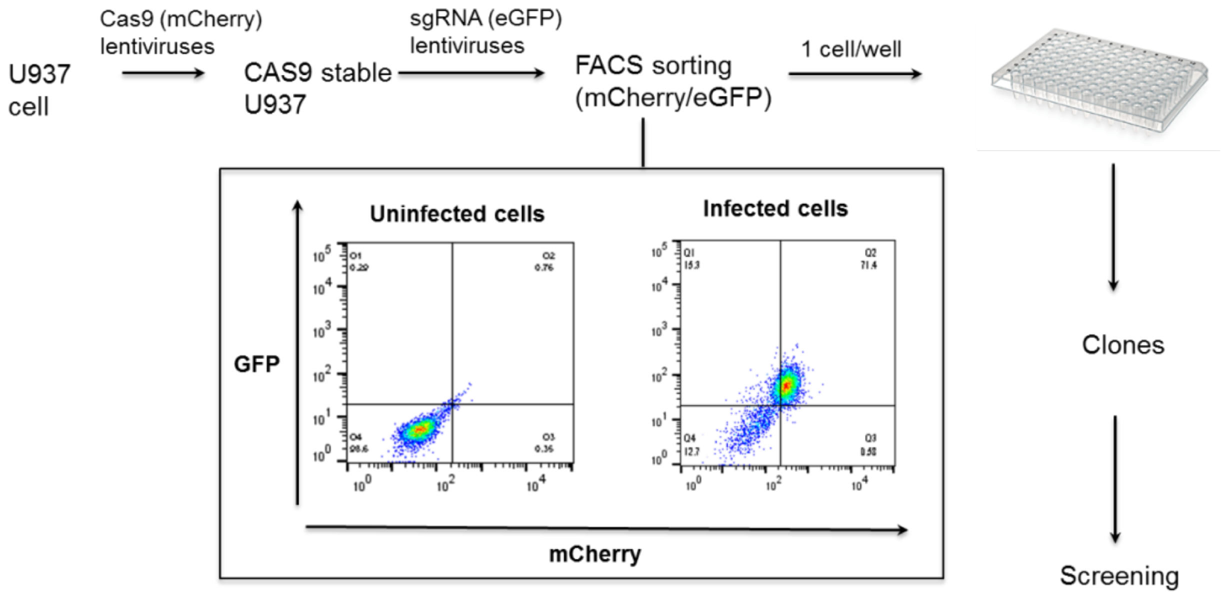


**Indel composition and frequency analyses by ICE**

**B**

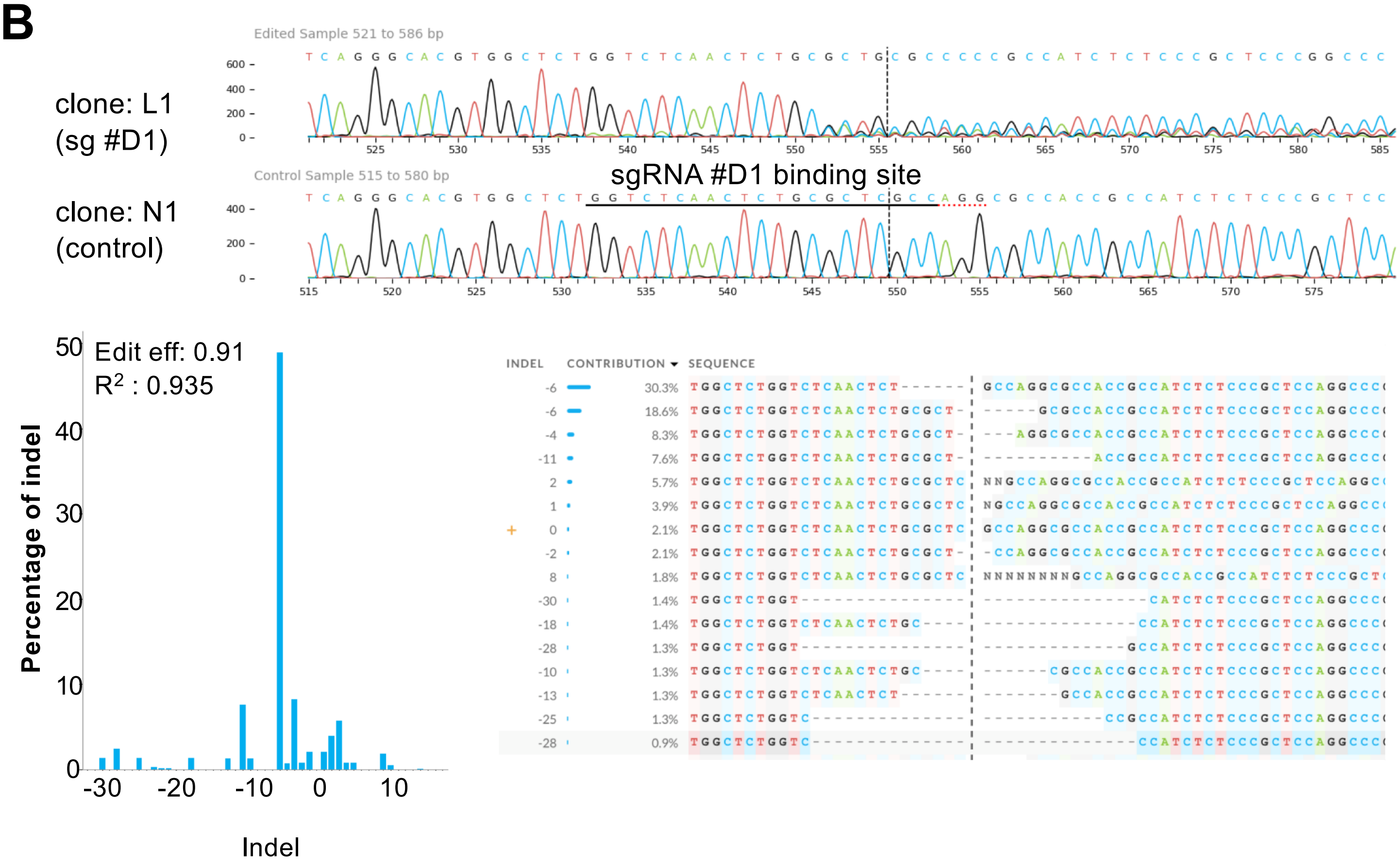


**FigS4**

**C**

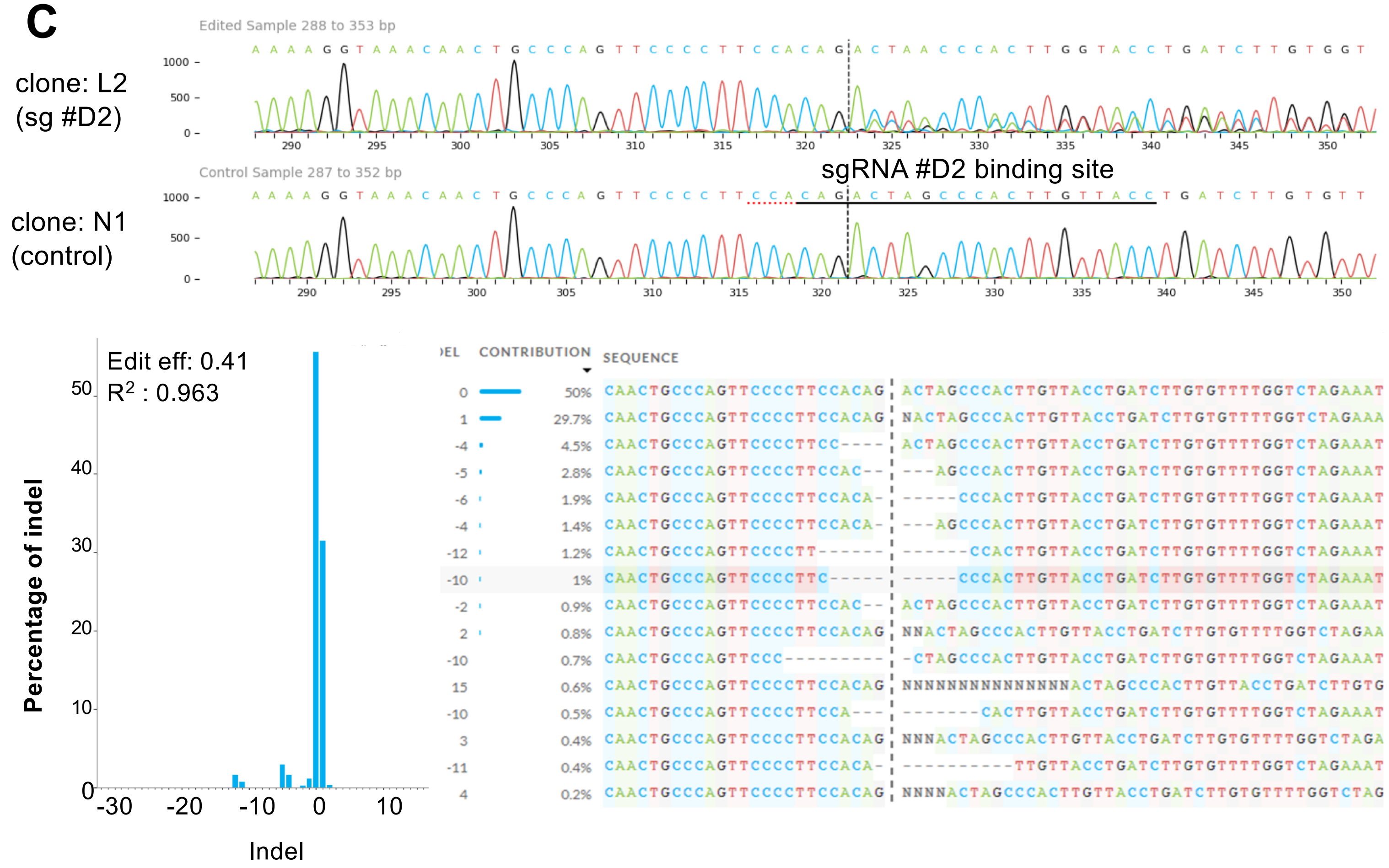


**D**

**Genomic DNA sequence at sg #D2 targeting region**

| | | |
|---|---|---|
| | clone N1 (control) | CTAGACCAAAACACAAGATCAGGTAACAAGTGGGCTAGTCTGTGGAAGGGGAACTGGGCAGTTGTTTACCTTT |
| Homologous indels | clone L2a (sg #D2) | CTAGACCAAAACACAAGATCA---------------------------GGGAACTGGGCAGTTGTTTACCTTT |
|  | clone L2b (sg #D2) | CTAGACCAAAACACAAGATCAGGTAACAAGTGGGCTAGT----GGAAGGGGAACTGGGCAGTTGTTTACCTTT |

Green: upstream plus gRNA binding site
Red: downstream of gRNA binding site

**FigS4**

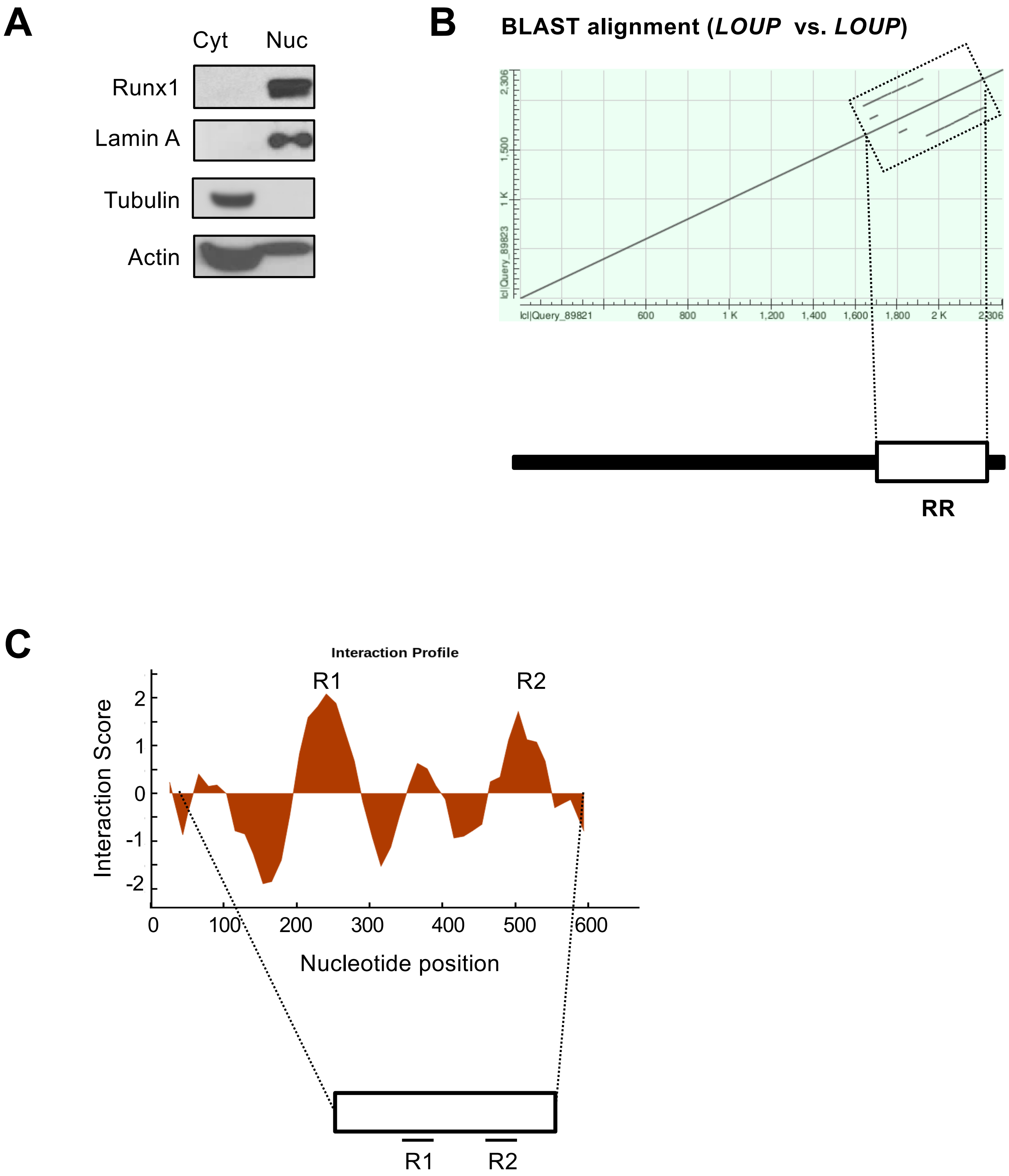
A
Cyt
Nuc
Runx1
Lamin A
Tubulin
Actin
B
BLAST alignment (LOUP vs. LOUP)
RR
C
Interaction Profile
R1
R2
Interaction Score
Nucleotide position
0
100
200
300
400
500
600
R1
R2

FigS5
